# Long-read sequencing of diagnosis and post-therapy medulloblastoma reveals complex rearrangement patterns and epigenetic signatures

**DOI:** 10.1101/2022.02.20.480758

**Authors:** Tobias Rausch, Rene Snajder, Adrien Leger, Milena Simovic, Oliver Stegle, Ewan Birney, Marc Jan Bonder, Aurelie Ernst, Jan O. Korbel

## Abstract

Cancer genomes harbor a broad spectrum of structural variants (SV) driving tumorigenesis, a relevant subset of which are likely to escape discovery in short reads. We employed Oxford Nanopore Technologies (ONT) sequencing in a paired diagnostic and post-therapy medulloblastoma to unravel the haplotype-resolved somatic genetic and epigenetic landscape. We assemble complex rearrangements and such associated with telomeric sequences, including a 1.55 Megabasepair chromothripsis event. We uncover a complex SV pattern termed ‘templated insertion thread’, characterized by short (mostly <1kb) insertions showing prevalent self-concatenation into highly amplified structures of up to 50kbp in size. Templated insertion threads occur in 3% of cancers, with a prevalence ranging to 74% in liposarcoma, and frequent colocalization with chromothripsis. We also perform long-read based methylome profiling and discover allele-specific methylation (ASM) effects, complex rearrangements exhibiting differential methylation, and differential promoter methylation in seven cancer-driver genes. Our study shows the potential of long-read sequencing in cancer.

**Graphical abstract:** **I)** We investigate a single patient with chromothriptic sonic hedgehog medulloblastoma (Li-Fraumeni syndrome), with tissue samples taken from blood, the primary tumor at diagnosis, and a post-treatment (relapse) tumor. **II)** Data on the three samples has been collected from four sources, 1) Illumina whole-genome, 2) Illumina transcriptome sequencing, 3) Illumina Infinium HumanMethylation450k, as well as 4) long-read whole-genome sequencing using Oxford Nanopore Technologies (ONT) sequencing. **III)** An integrative analysis combines genomic, epigenomic as well as transcriptomic data to provide a comprehensive analysis of this heavily rearranged tumor sample. Long and short read sequencing data is used to inform the analysis of complex structural genomic variants and methylation called from haplotyped ONT reads and validated through the methylation array data allows for a haplotype-resolved study of genomic and epigenomic variation, which can then be examined for transcriptional effect. **IV)** This integrative analysis allows us to identify a large number of inter- and intra-chromosomal genomic rearrangements **(A)** including a complex rearrangement pattern we term templated insertion threads **(B)**, as well as sample-specific and haplotype specific methylation patterns of known cancer genes **(C)**.

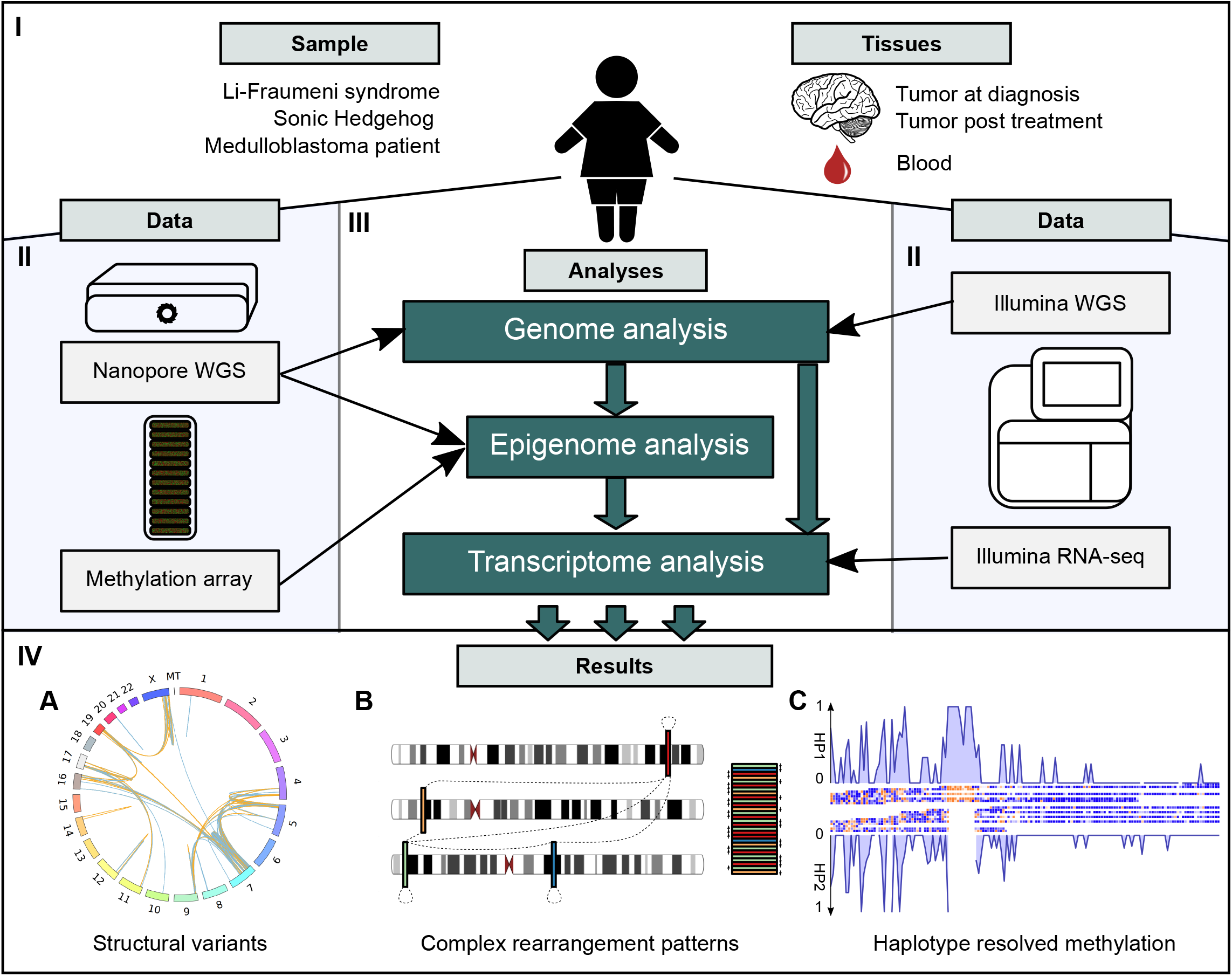

## Introduction

Cancer genomic landscapes are shaped by a diversity of somatic rearrangement patterns, ranging from simple deletions, duplications and reciprocal translocations to SVs formed via complex DNA rearrangements, including breakage-fusion-bridge cycles and chromothripsis events^1–4^. SVs are the most common source of cancer driver mutation, outnumbering point mutations for the generation of cancer drivers in the majority of common cancers^2^; yet, owing to technical difficulties with respect to their discovery and characterization^5^, their structure and patterns are underexplored compared to point mutations^2^. This is particularly true for complex DNA rearrangements, the characterisation of which remains an important challenge, with short-read (Illumina) sequencing data only partially resolving such structures^3^.

Initial efforts to classify somatic SVs uncovered a variety of common somatic rearrangement patterns, which suggest that a wide variety of rearrangement processes are active in cancer. Using non-negative matrix factorization, Nik-Zainal et al.^6^ initially described six signatures of rearrangement in breast cancers sequenced using Illumina technology. More recent pan-cancer studies^3,7^, again pursued using short read data, combined simple SVs (e.g. deletion-type, duplication-type and inversion-type) into discrete higher level patterns based on breakpoint junction connectivity, resulting in over a dozen SV signatures. This included patterns of intermediate rearrangement complexity, such as templated insertion chains comprising up to 10 breakpoints. Yet, more complex rearrangement patterns have so far largely resisted systematic classification based on breakpoint junction connectivity. An important reason for this is difficulty in assembling short-read data into coherent structural segments to study patterns of somatic rearrangements. This problem is exacerbated by repetitive sequences in the genome, in which SV breakpoints are readily missed by Illumina whole genomes sequencing (WGS). This leaves open the possibility that important patterns of structural rearrangement have not yet been discovered and are elusive due to the predominant use of short-read sequencing in cancer genomics^2^.

Here we sought to evaluate the utility of long read sequencing technology^8–11^, in particular Oxford Nanopore technology (ONT), to reveal patterns of somatic structural variation. The technological choice was motivated by the fact that long read sequencing of 1000 Genomes Project samples showed a greatly increased number of confidently discovered SVs in repetitive regions, improved sensitivity for SVs smaller than 1 kbp in size, and advantages for investigating complex SV patterns by facilitating haplotype-resolved genomic sequence assembly^12,13^. ONT additionally shows great promise in cancer epigenomics, as from the same long reads both genetic and DNA methylome data can be obtained, the latter of which is quantified through measuring current changes within the nanopore^14^ – which should allow integrated characterization of genetic and epigenetic changes in tumors at single (long) molecule level. However, there is a current lack in suitable computational methods and hence a need in exploring and devising approaches leveraging long read data in cancer genomes – with the complications of intra-tumor heterogeneity in primary cancer samples, normal cell contamination, aneuploidy and complex SVs, and variation in tumor methylation levels.

To address the current lack of long-read analytical methods to explore cancer genomes, we performed ONT sequencing of a childhood medulloblastoma, and devised methods to enable characterizing SV and methylome patterns in these data. The tumor arose in a patient carrying a germline *TP53* mutation (Li-Fraumeni syndrome, OMIM Entry # 151623), previously associated with Sonic-Hedgehog subgroup medulloblastoma (SHH-MB) and somatic chromothripsis^15,16^. We reveal the fully assembled haplotype-resolved structure of a complex chromothripsis event^15,17^. We further uncover a novel complex rearrangement pattern, termed templated insertion thread, which copies and concatenates a substantial number of short subkilobase-sized templated insertions in forward and reverse orientation, resulting in massively amplified sequences ranging up to several tens of kilobases in size. While not initially discovered by Illumina WGS, we demonstrate that common features associated with templated insertion threads allow their discovery in cancer genomes sequenced with short-reads. A search for these patterns in 2,569 short read cancer genomes from the Pan-Cancer Analysis of Whole Genomes (PCAWG) consortium^2^ reveals templated insertion thread footprints in 3% of cancer genomes, with a particular abundance in liposarcoma (74%), glioblastoma (24%), osteosarcoma (22%) and melanoma (14%). Templated insertion threads can occasionally be linked to cancer-related gene overexpression, suggesting that cancer cells could exploit this somatic SV pattern to promote tumor evolution. Lastly, by integrating genomic and epigenomic readouts, we performed haplotype-resolved genome-wide analysis of CpG methylation. We associate a subset of the somatic DNA rearrangements, including templated insertion threads, with functional consequences, and demonstrate the ability to explain aberrant gene expression patterns, such as allele specific expression and gene-fusions, by integrating genomic and epigenetic long read data.

## Results

### ONT-based integrated phasing and SV discovery in a medulloblastoma patient

We sequenced the primary medulloblastoma (sample ID: LFS_MB_P) to ~30x ONT coverage, and generated ~15x for a tumor specimen taken during relapse (LFS_MB_1R) and a paired blood control sample, respectively, with a median read length of 5kbp (**Table S1**). We developed workflows and algorithms to analyze both genetic and epigenetic alterations in these samples (**Methods**). Making use of short-read data generated at 45x-48x coverage for these sample^s16,18,19^ (**Table S2**), we discovered single-nucleotide variants (SNVs) as well as short insertions and deletions (InDels), where ONT reads have limitations due to their relatively high error rate. As expected, germline variant calling confirmed a *TP53* mutation (TP53:c.395A>T, p.Lys132Met, rs1057519996), consistent with Li-Fraumeni Syndrome, coupled with somatic inactivation of the wild-type *TP53* allele through deletion in the tumor samples. To facilitate allele-specific analysis we devised a haplotype-phasing approach that generates initial haplotype blocks from ONT reads, which then are integrated with statistical haplotype phasing data from the 1000 Genomes Project^20^; haplotype switch errors are then corrected by leveraging somatic copy-number alterations (SCNA) in the tumor that result in allelic shifts away from the normal 1:1 haplotype ratio (**Figure S2**). In regions of the genome without SCNAs we estimate an N50 phased block length of 4.68 Mbp using this approach (**Methods**). The estimated proportion of the somatic genome that is haplotype-resolvable using our phased germline variant call set is 93.6% for the primary tumor and 90.9% for the relapse sample, respectively.

### Haplotype-phased assembly of complex somatic rearrangements

We integrated ONT-based somatic SV calling with Illumina-based SCNAs and variant detection to achieve haplotype-resolved reconstruction of the somatic SV landscape of this tumor (**Methods**). In the primary tumor, we find 697 somatic SVs, including 106 deletion-type SVs, 107 duplication-type SVs, 189 inversion-type SVs and 295 inter-chromosomal rearrangements. Most of these rearrangements arose from two distinct chromothripsis events – one involving chromosomes 4, 5, 7, 9, 16, 19 and X, and the other chromosomes 11 and 17, respectively (**Figure 1AB**, **Figure S4**). We explored targeted phased assembly of the genomic outcomes of both chromothripsis events (**Methods**), and constructed SV contigs for the chromothripsis event spanning chromosomes 4, 5, 7, 9, 16, 19 and X, and a phased assembly of fragments originating from chromosome 11 and 17 (denoted CS11-17, **Figure 1CD**). The CS11-17 segment, present in both primary tumor and relapse, has a size of 1.55 Mbp; the 17p-arm region affected contains the *TP53* locus, which has been lost on the chromothriptic haplotype. We estimate an average copynumber of 3 to 4 copies for CS11-17, consistent with FISH experiments (**Table S3**). FISH further shows extensive intra-tumor heterogeneity (ITH) of CS11-17 copy-numbers, which range from 1 to 7 **(Tables S3**, **S4**, **S5**). We performed sequence-level characterization of CS11-17, and partially resolved peri-centromeric regions at its flanks (**Figure 1CD**), which could provide the necessary sequence context for homology based integration into the normal genome as observed previously for double minutes^17^ (**Figure 1E**). Indeed, the absence of classical double-minute chromosome structures in metaphase spreads analyzed by FISH suggests the likely reintegration of CS11-17 (**Figure S5**). Yet, we failed to identify reads supporting reintegration of this structure into a chromosomal context, possibly due to limitations of ONT for resolving low-variant allele frequency SVs in conjunction with ITH, especially in complex regions that exhibit repetitive segments larger than the ONT read length^21^.

**Figure 1.**
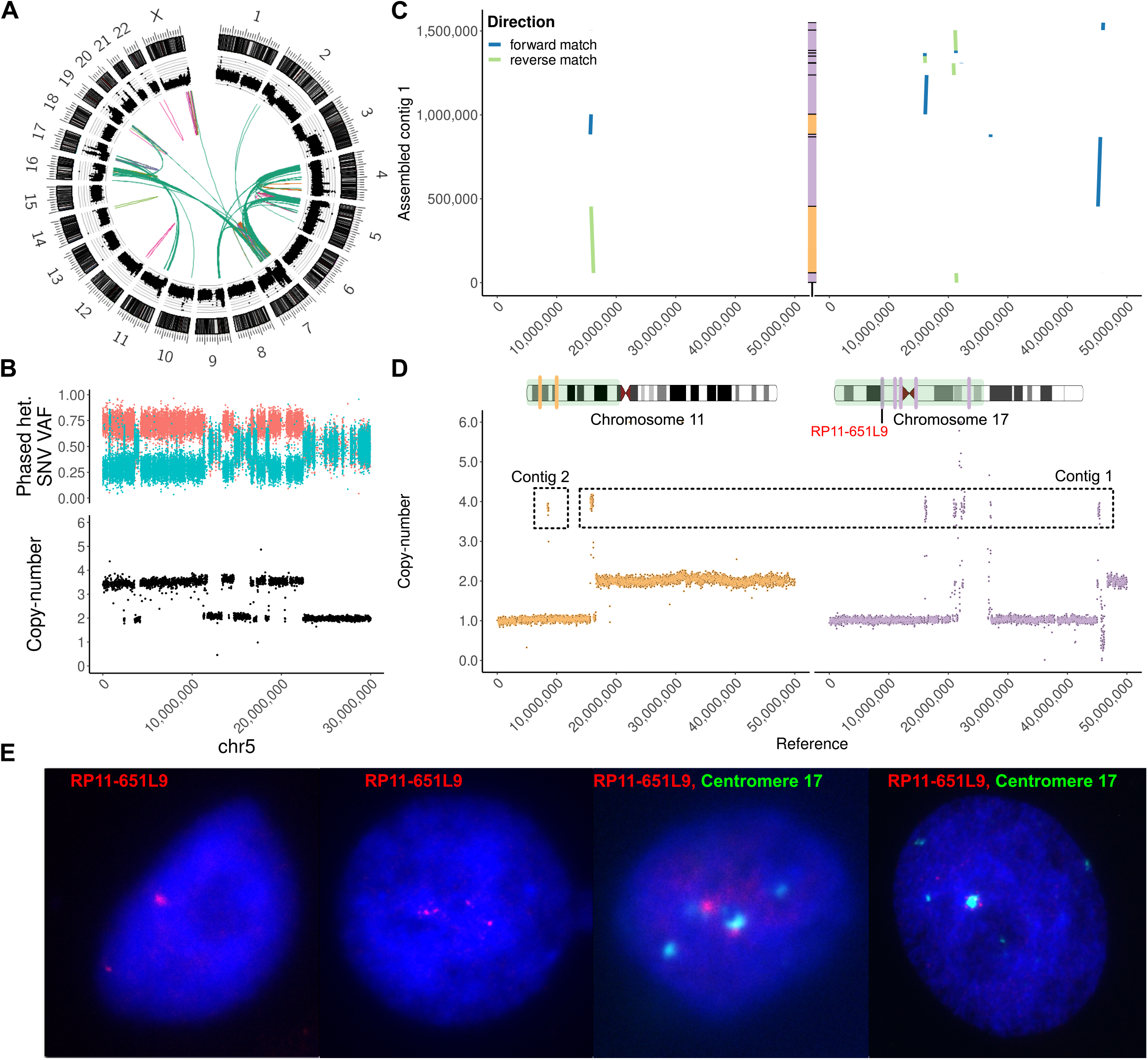
Haplotype-phased assembly of an inter-chromosomal chromothripsis event. **(A)** A circos plot of the primary tumor showing from outside to inside the chromosome ideograms, read-depth, large (>10Mbp) structural variants and inter-chromosomal rearrangements with orange: deletion-type, violet: duplication-type, light-green: head-to-head inversion-type, pink: tail-to-tail inversion-type and dark-green: inter-chromosomal. **(B)** Chromosome 5 exhibits a pattern of oscillating copy-number states (lower panel) and alternating heterozygous allele frequencies (upper panel) common to chromothripsis. **(C, D)** The CS11-17 assembly is aligned to chromosome 11 and chromosome 17 with aligned segments corresponding to amplified regions at approximately copy-number 4 in panel D. Segments from chromosome 11 are in yellow, segments from chromosome 17 in purple. The subset of the chromosomes displayed (1-50Mbp) is highlighted in green in the chromosome ideograms as well as the location of the amplified segments. **(E)** FISH pictures of the red RP11-651L9 probe (chr17:16,169,409-16,359,715) and the green centromere 17 probe showing distinctive intra-tumor heterogeneity for the CS11-17 structure. From left to right, (i) nucleus showing 2 signals for the RP11-651L9 probe, (ii) 4 signals for the RP11-651L9 probe, (iii) colocalization of the centromere 17 probe with the RP11-651L9 probe, and (iv) clusters of signals for the RP11-651L9 probe around the centromere 17, suggesting a possible peri-centromeric integration.

### ONT sequencing reveals a novel complex rearrangement pattern denoted templated insertion thread

Notably, the somatic SVs included a highly unusual pattern of inter-chromosomal DNA rearrangement not matching previously described somatic SV classes. This rearrangement pattern involves short DNA segments, mostly 100bp–1kbp in size, that are concatenated by a structural rearrangement process in forward and reverse order, into a complex, highly amplified sequence comprising up to 50kbp of DNA and dozens to hundreds of breakpoint junctions (**Figure 2A**). We find two such structures in the primary tumor, yet, identify no such pattern in the relapse sample. We analyzed this unusual rearrangement pattern more closely and found that the length of the source sequence segments ranges from 144 - 3,637 bp, with all source segments with an estimated total copy-number greater than 10 being between 225 bp and 403 bp in size. The total length of the resulting somatic amplicon structure is 50.3kbp for the first structure (**Figure 2B**) and 39.9kbp for the second structure (**Figure S6**). Both of these structures result in inter-chromosomal adjacencies, via concatenation of templated insertions stemming from distinct chromosomes. Self-alignments of ONT reads spanning the amplicon structure independently verified the repetitive nature of these insertions (**Figure 2AB**, **Figure S6, S7**). Based on a sequence analysis of these structures, and leveraging the full length of the ONT reads, we find that these structures most likely emerge from templated insertions^3^, which through a copy-and-paste process become concatenated in forward and reverse orientation with no apparent regularity with respect to the orientation of the concatenated source sequence segments (**Figure 2C, S8**) – and we therefore term this novel pattern ‘templated insertion thread’.

**Figure 2.**
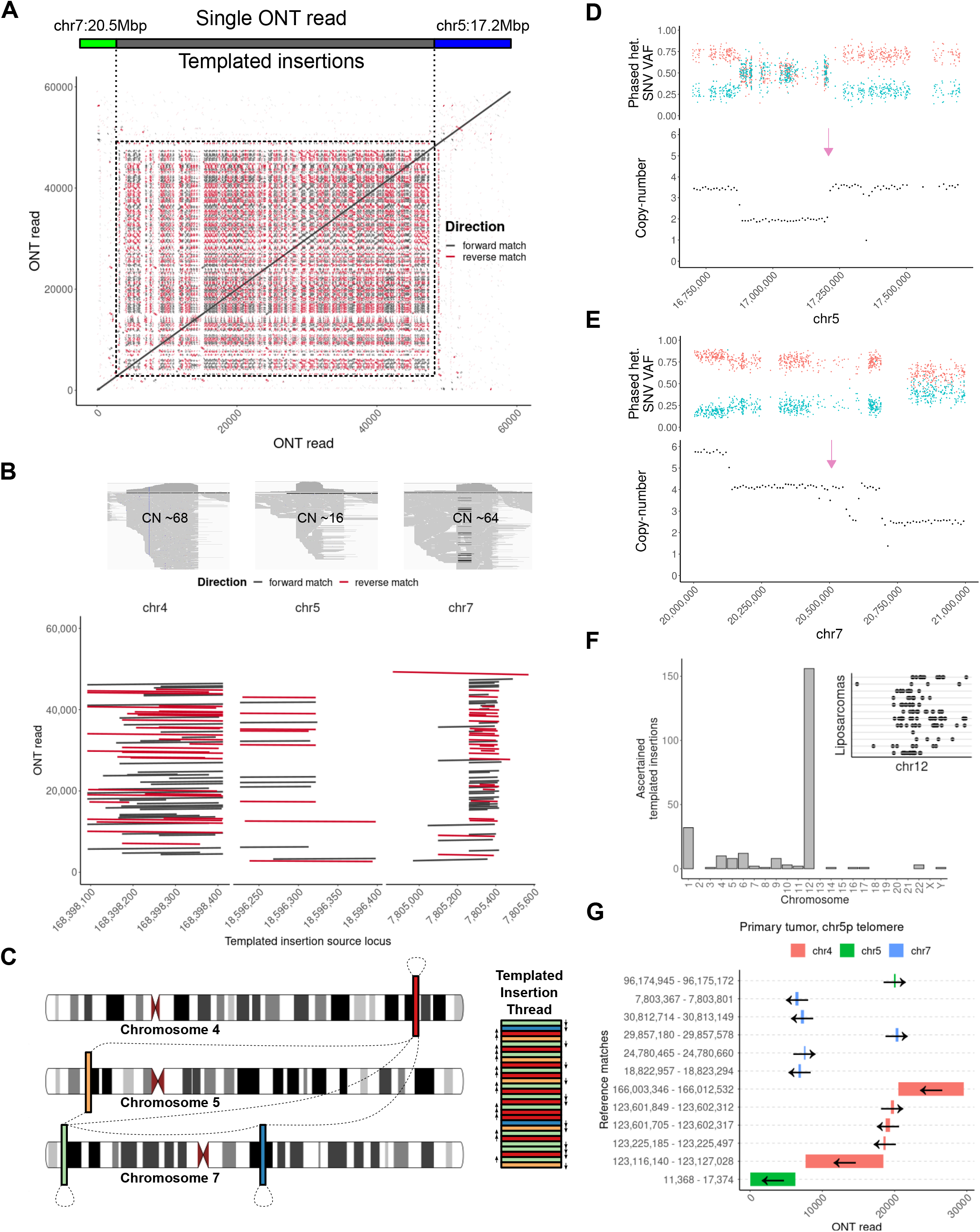
Templated insertion threads. **(A)** Self-alignment of a single ONT read that spans the entire length of the templated insertion thread, displaying an array of repetitive short sequence matches reflecting the copying and concatenation of few source sequence segments. **(B)** Matched illumina data shows a characteristic coverage increase (upper panel). An alignment of the ONT read (y-axis) against selected templated insertion source sequences (x-axis) shows how the ONT read aligns across these source sequences multiple times in seemingly random order. **(C)** A scheme showing how templated insertions are copy and paste in direct adjacency and random order into a growing templated insertion thread. Arrows next to the templated insertion thread indicate the segment orientation and dashed lines show discovered adjacencies among individual templated insertions. **(D)** The colocalization of the beginning and the end of the templated insertion thread (purple arrow) with chromothripsis segments on chromosome 5 and **(E)** chromosome 7. **(F)** Analysis of 2,569 cancer genomes reveals that liposarcomas often harbor templated insertion threads, preferentially on chromosome 12 (main panel). The inset shows the distribution of templated insertions along chromosome 12 where each horizontal line is a distinct liposarcoma sample. **(G)** Telomeric repeat analyses identified a complex SV rearrangement involving chromosome 4, 5 and 7 that was stabilized by telomere fusion to the chr5p telomere in the primary medulloblastoma sample.

A comparison with previously described rearrangement patterns shows that the templated insertion thread pattern shares features with the chains of templated insertions pattern previously described by Li et al. using PCAWG data^3^ and the tandem short template jumps signature previously uncovered by Umbreit et al. in cell cultures^22^ – albeit with clear differences. While all these patterns concatenate templated insertions originating from distinct genomic locations, the most distinguishing feature of templated insertion threads is the prevalent self-concatenation of templated insertions in a zig-zag fashion, which result in short amplicons of remarkably high copynumber (**Figure 2BC, S9**); by comparison the units comprising chains of templated insertions occur only once (no self-concatenation) in the previously described patterns^3,22^. As an additional discriminating feature, chains of templated insertions as described by Li et al.^3^ comprise from 1 to 10 concatenated units, compared to >50 units included within a single templated insertion thread in this medulloblastoma sample (see **Figure S9**).

We performed further analyses of the spanning ONT reads, and found that the templated insertion threads colocalize with chromothriptic rearrangements (**Figure 2DE**). It is therefore possible that the rearrangement processes resulting in both event classes share some commonality, either with one event triggering the other, or with both chromothripsis and templated insertion threads enabled by the same initiating DNA lesion. Analysis of the repeat units (source sequence segments) becoming self-concatenated in templated insertion threads did not reveal any biases towards a specific sequence context; in the majority of cases, individual units originate from non-repetitive sequence (**Methods**). Interestingly, comparative alignment of ONT reads from the same sample revealed evidence for ITH with respect to the unit composition of templated insertion threads, with clear differences in concatenated unit numbers becoming evident; this suggests that sites of templated insertion thread events may be prone to undergo further somatic rearrangements generating further genetic heterogeneity **(Figure S10)**.

### Graph-based discovery of templated insertion threads in Illumina WGS data

Most previously sequenced cancer genomes have been generated using short reads, which compared to long reads display poor sensitivity towards <1kb-sized rearrangements^13^ – the predominant rearrangement type within templated insertion threads. Irrespective of this, we hypothesized that the distinguishing features of templated insertion threads should be discoverable in short read data once explicitly sought for – to allow further analysis of this novel SV pattern in large short-read based cancer genome cohorts. To address this hypothesis, we first closely examined the Illumina WGS reads from LFS_MB_P at the sites of templated insertion threads. Indeed, we find specific short read alignment patterns characteristic of self- and cross-linked sequence segments at the respective rearranged sites, with exceptionally high copy-number of source segments and paired-end as well as split-read support for rearrangement junctions **(Figure S11)**. Encouraged by this observation, we devised the graph-based algorithm *rayas*, to enable the discovery and characterization of templated insertion threads in short read WGS data (**Methods**). The algorithm combines read-depth and split-read patterns to identify rearrangement graphs, allowing the specification of 1:n relationships, whereby a single templated insertion source sequence (i.e., a node in the graph) can contribute to different rearrangement adjacencies (i.e., edges in the graph; **Figures S11**). Application of *rayas* to the primary and relapse samples led to the re-discovery of both templated insertion threads in the primary medulloblastoma, and confirmed the absence of these structures in the relapsed medulloblastoma.

### Pan-cancer landscape of templated insertion threads in 2,569 tumors

The ability of template insertion threads to amplify short sequences suggests a potentially broader relevance in cancer, since amplified DNA sequences could potentially act as cancer drivers such as by focally amplifying DNA regulatory sequences or altering the gene regulatory context to result in ectopic expression^2,23,24^. To enable a wider characterization of this SV pattern, we used *rayas* to interrogate 2,569 cancer genomes from the PCAWG consortium^2^. We find 169 templated insertion threads in 76 (~3%) cancer genomes, which suggests that this somatic rearrangement pattern arises in distinct cancers (**Figure S12, Table S6**). Across cancers the distribution of this pattern is highly heterogeneous, with 74% of liposarcomas, 24% of glioblastoma and 14% of melanomas exhibiting template insertion threads, versus 7% of leiomyosarcomas **(Figure S12)**. We caution that due to the lower sensitivity of short-reads for detecting complex SVs involving short repeat units^13^, future studies with larger cohorts of cancer samples sequenced with long-reads will likely reveal a higher frequency of templated insertion threads in cancer.

On average, templated insertion threads consist of 4 distinct source segments with a median unit size of 558bp, and median number of concatenated units of 53.1, indicating that high copy number amplification is the norm rather than the exception for this SV pattern. We next analyzed these 76 cancer genomes bearing template insertion threads in more detail, to determine features that may potentially correlate with the occurrence of template insertion threads. Interestingly, 65 out of these 76 cancer genomes (86%) were previously classified as having at least one chromothripsis event^2^. The association of template insertion threads with chromothripsis is significant across 2,569 cancers, when adjusting for tumor histology, gender and ancestry (p-value: 1.15 × 10^−5^, logistic regression). Interestingly we find a strong enrichment of templated insertions on chromosome 12 in liposarcoma samples, with a propensity towards the 12q15 chromosome band (**Figure 2F**). Liposarcomas often form supernumerary ring or giant marker chromosomes that include multiple copies of the target oncogenes (*MDM2, CDK4*, among others) on chromosome 12, a chromosome that frequently undergoes chromothripsis in this cancer type^18,25,26^. A recent study also identified chromosome 12 as a hotspot for seismic amplification in liposarcoma^27^. These data suggest that templated insertion threads could arise in association with supernumerary ring or giant marker chromosomes, possibly triggered by the same initiating lesions or through a common rearrangement process.

### Telomere analysis of derivative chromosomal segments

Critical telomere shortening is one mechanism implicated in triggering complex structural rearrangements such as chromothripsis events^28,29^. Prompted by complex inter-chromosomal rearrangement seen in this medulloblastoma patient, we explored telomeric sequences associated with the resulting derivative chromosome structures, an analysis normally inaccessible to short reads. We devised a method to identify telomeric motifs, repeats of TTAGGG, TGAGGG, TCAGGG, TTGGGG or their reverse complement, in error-prone ONT reads and applied this method to the long read data of the primary tumor and the relapse sample (**Methods**). Using this approach, we confidently detect five structural rearrangements involving telomeric sequences – three in the primary tumor and two in relapse – where a telomeric sequence of one chromosome is fused to a rearranged segment of another chromosome (**Figure 2G**, **S13**). For one of these telomeres we identify a highly complex rearrangement pattern, involving the chromosome 5p telomere and several short sequence segments from chromosome 4, 5, and 7 (**Figure 2G**) reminiscent of chains of templated insertions. For this event, telomere crisis may have initiated the complex SV pattern present throughout chromosome 4, 5 and 7, including chromothripsis and the above mentioned templated insertion threads. Telomere fusions can also stabilize altered chromosomes after catastrophic events such as chromothripsis^30^, which would suggest an alternative sequence of events, with chromothripsis and templated insertion threads causing unprotected break sites healed through telomere addition. Another telomere crisis event observed in the primary tumor likely fused chromosome 19 to the telomere of chromosome 16q, an event that could only be resolved unambiguously using the CHM13 telomere-to-telomere (CHM T2T) assembly^31^ as a reference sequence (**Figure S13**). We further investigated whether eroded telomeres were preferentially fused with genomic loci active in transcription as has been suggested previously^32^, but our small number of telomere fusions do not provide sufficient evidence. Telomeres can erode more rapidly in cells of Li-Fraumeni syndrome patients as compared to healthy individuals, which is thought to lead to an increased frequency of telomeric fusions^33^, and possibly contributed to the complex SV patterns observed in this study.

### Differential methylation from long-read data

ONT sequencing allows for direct assessment of the methylation likelihood of cytosine bases^14^, providing the opportunity to characterize global DNA methylation levels in this medulloblastoma sample, and to integrate DNA methylome and somatic rearrangement data. We quantified DNA methylation at base-level resolution using Nanopolish, which yields good correlation (pearson-R^2^ 0.9102 in primary tumor, 0.8497 in relapse) with methylation rates obtained through the HumanMethylation450 array platform (**Figure S14**).

We attempted to identify patterns of variation in DNA methylation by comparing methylation rates between primary tumor and relapse sample using PycoMeth^34^. We find that directly testing methylation rates of gene promoter regions (as defined in methods) yields poor power, with only 31 gene promoters called as differentially methylated (FDR <= 0.05, abs methylation rate difference > 0.5). We therefore apply two segmentation approaches, testing for differential methylation in segments defined using PycoMeth’s CGI finder and PycoMeth’s *de novo* methylome segmentation method Meth_Seg respectively (**Methods**). The between sample segmentation identified 662,262 methylation-based segments as well as 358,922 CpG-dense regions. Differential methylation calling on the segmented methylation calls reveal 2,459 individual segments, or 26,542 CpG sites, called as differentially methylated (**Figure 3A**) with an average length of 402 base pairs per segment (FDR <= 0.05, abs methylation rate difference > 0.5, **Figure S15**). Of these CpG sites, 3,117 (11.74%) intersect with gene promoters, revealing 475 genes with differential promoter methylation, seven of which were previously annotated as medulloblastoma driver genes^35^ representing a significant enrichment (Fisher’s exact test statistic: 20.25, p-value: 1.6 × 10^−7^). Furthermore, 742 (2.80%) CpG sites intersect with 64 enhancers active in the cerebellum. Among these we detect hypermethylation in an enhancer and promoter region of the neuritin 1 gene (*NRN1*) (**Figure 3B**), previously identified as down-regulated in treatment-resistant medulloblastoma^36^ and linked with tumor growth suppressive features in esophageal cancer^37^. We also observe a 329bp region in the promoter of *PTCH1*, a key driver in Sonic Hedgehog medulloblastoma^38^, which is methylated in the relapsed tumor and heterozygously deleted in both samples. Overall, analysis of the ONT data provides a substantially more comprehensive picture of the tumor methylome, with 78% of the between sample DMRs inaccessible to the commonly used 450K array, and 65% inaccessible to the 850K array **(Figure S16**).

**Figure 3.**
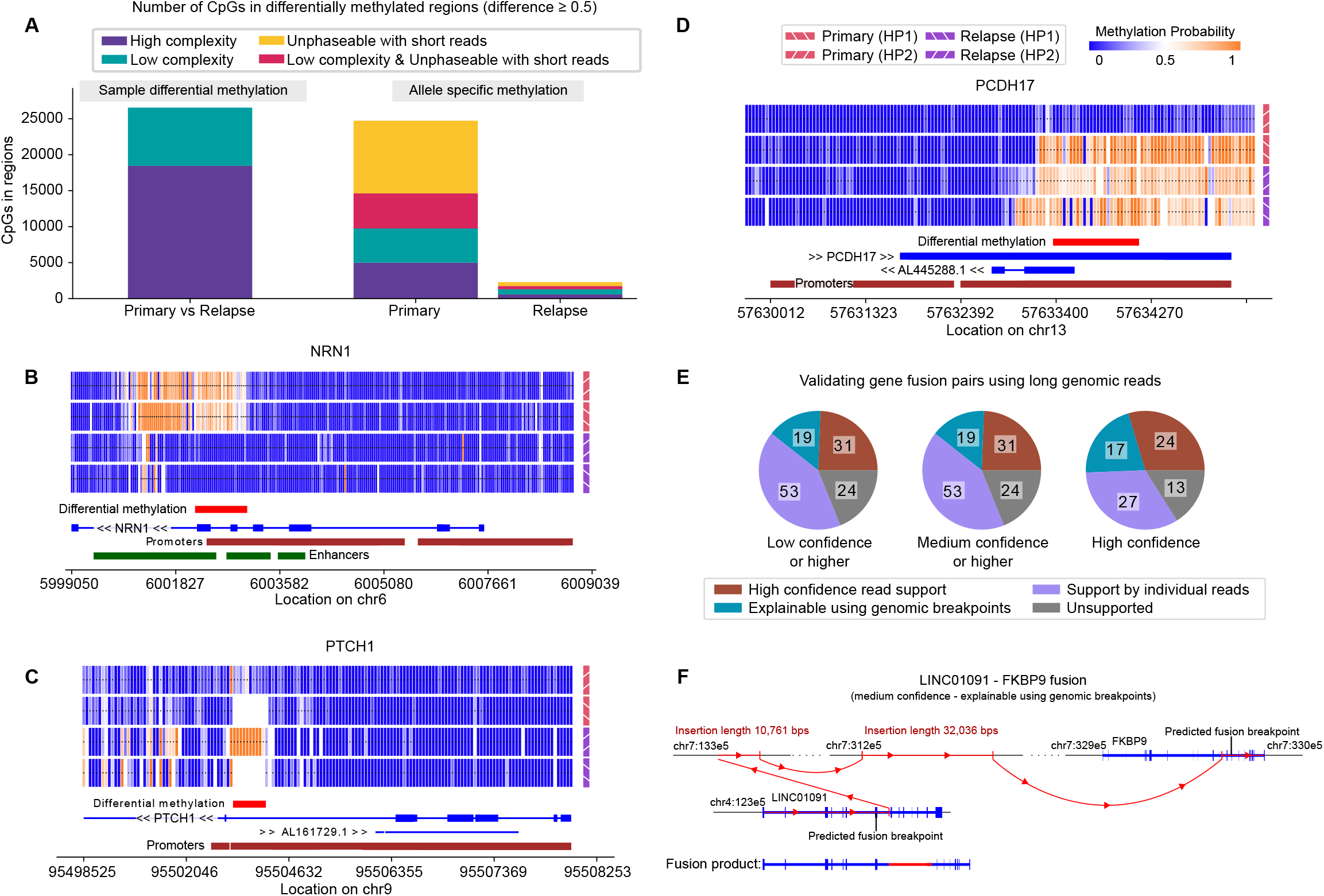
Functional analysis of primary tumor and relapse sample. **(A)** Number of CpGs in regions found to be differentially methylated in the sample comparison (Primary tumor vs Relapse) as well as ASM in the two samples. Colors represent an estimation of discoverability with short-read sequencing methods. CpGs in low complexity regions (soft-masked in reference) are more difficult to map using only short reads. CpGs not phaseable with short reads are further than 150bps from a phased heterozygous non C>T variants. **(B)** Methylation of *NRN1* promoter and enhancer in the primary tumor sample. **(C)** Heterozygous deletion in promoter of *PTCH1* (tumor suppressor gene and driver in Medulloblastoma) with differential methylation in the remaining haplotype. **(D)** *PCDH17* (tumor suppressor gene) promoter with ASM pattern in the primary tumor sample. **(E)** Predicted gene fusion pairs from Arriba validated using ONT long read information, thresholded by confidence as reported by Arriba. Fusion pairs in the *Supported by individual reads* category are supported by at least one genomic read with a chimeric alignment including both genes. Pairs in the *Explainable using genomic breakpoints* category have a plausible explanation by following a graph of structural variations that connect the two genes. The category *High confidence read support* refers to pairs where both these criteria are met. **(F)** Example of a gene fusion pair that can be explained using genomic breakpoints but with no individual genomic read that covers both genes. Two separate insertions of a total length of 42,797 base pairs appear to be involved in the fusion of *LINC01091* and *FKBP9* such that even in ONT reads there was no read extending across the entire gene fusion.

### Resolving expression effects using ONT data

Leveraging Illumina RNA sequencing data generated for both primary tumor and relapse, we assessed whether differential methylation measured in gene promoters is associated with expression changes. Gene expression analysis revealed 49 genes with strong differential expression between the two samples (absolute log fold change >5 (a-l2fc), methods, **Table S7**), including in known medulloblastoma genes (amongst others *KCNA1*, and *DMBT1*)^39,40^. Of the total 475 promoter linked DMRs (415 are expressed in both samples), 57 overlap with differentially expressed genes (a-l2fc >2); the overlap between differential expression and DMR effects is statistically significant (Fisher’s exact test statistic: 12.27, p-value: 4.3 × 10^−6^). As previously described promoter methylation has a mostly negative relation to expression^41^, 50 out of the 57 pairs (87.7%), are negatively correlated, and we observe a significant inverse correlation (Spearman R: −0.31, p-value: 1.8 × 10^−2^) between methylation and expression levels (**Figure S19**). For example, we find that the *BCAT1* gene is overexpressed and under-methylated in the relapse, consistent with a prior report observing that this gene is overexpressed in metastatic compared to non-metastatic medulloblastomas^42^. We also find *TBX1* which is regulated by Sonic Hedgehog^43^ with two separate promoter-linked DMRs, one hypermethylated and one hypomethylated in primary tumor, while underexpressed (5.29 l2fc) in the primary tumor as compared to the relapsed tumor (**Figure S20**).

We further sought to integrate the transcriptomic data with the long ONT reads to look for supporting data for gene fusion events (see **Table S7**), previously described to be prevalent in SHH-Medulloblastoma^44^. We inferred gene fusion events from transcriptomic reads using Arriba on the primary tumor, and identified 127 putative gene fusion pairs of which 103 pairs are supported by genomic evidence, either directly through individual chimeric read alignments of ONT reads near the fusion breakpoints (53) or by tracing SVs called from long and short genomic reads (19) or both (31) (**Methods**). Breaking down predictions by Arriba confidence shows increased traceability for higher confidence fusion calls (**Figure 3F**). Tracing SVs, across a limited number of ONT reads, allows us to explain long and complex fusions, such as the gene fusion observed between *FKBP9* and *LINC01091*, with the fusion breakpoint in a long (>69kbps) intron resulting in an intronic insertion of 42,797bps length (**Figure 3G**). Interestingly we observe a translocation involving *NCOR1* and *AC087379.1*, genes on the CS11-17 structure. *NCOR1*, a tumor suppressor gene, has previously been reported in loss-of-function fusions in SHH medulloblastoma^44^; the *NCOR1-AC087379.1* fusion detected here is out of frame and therefore would be predicted to disrupt *NCOR1*.

### Allele specific methylation and expression

ONT sequencing gives the unique opportunity to phase long methylation called reads, allowing high resolution allele specific methylation (ASM) analyses along the cancer genome. We analyzed ASM patterns, by running a second segmentation using PycoMeth Meth_Seg, a methylome segmentation method leveraging sample haplotype information (**Methods**). Using the same FDR cutoff as for DMR analysis (**Methods**), we identify 1,171 differentially methylated segments between the haplotypes of the primary tumor sample, spanning a total of 24,725 CpGs, with an average segment length of 525 base pairs (**Figure S16**). Due to the lower sequencing depth in the relapse sample, the number of segments passing the significance threshold with ASM is lower, resulting in 77 differentially methylated segments (spanning 2,289 CpGs, **Figure 3A**). While detection power in relapse is low due to lower read-depth, 401 of the 1,172 ASM segments (34.22%) found in the primary tumor show the same effect in the relapse sample with regards to sign and methylation rate difference (**Methods**). To illustrate the benefit of using non bisulphite converted long reads for this analysis we separate out CpGs close to heterozygous variants (<=150bps away) versus CpGs further away from heterozygous variants (excluding C>T variants as those cannot be distinguished from methylation calls in bisulfite sequencing) observing that we can get 19,729 (395%) more CpGs confidently linked to ASM effects (**Figure 3A**).

In the primary tumor sample, a total of 396 gene promoters and 29 enhancers intersected with segments with ASM, and 23 gene promoters and 1 enhancer in the relapse sample. Among these, we observe promoter methylation of *PCDH17*, a tumor-suppressor gene in which aberrant promoter methylation was previously observed in different tumors^45–49^. We also detected longer segments, such as a 26,751bp long region found as part of a larger ~250kbp long region on chromosome 15 spanning three protein coding genes as well as a 53 non-coding genes including the *SNORD116* and *SNORD115* clusters, which is partially methylated in one haplotype and fully methylated in the other. The full list of genes with sample specific or allele specific methylation can be found in **Table S8**. Unable to confirm a significant relationship between ASM and proximity to somatic variants, it is likely that a sizable fraction of ASM detected is associated with germline variation.

We also investigated whether ASM is associated with gene expression levels, by performing allele specific expression analysis. Using the phased variants from the blood sample, we are able to compute ASE rates using WASP (**Methods**), focusing on the variants in the gene promoter region as defined for ASM. We observe a total of 220 genes with a significant ASE effect (Q-value <0.05). A total of 70 genes that show ASE effects were previously implicated in medulloblastoma, including the previously described *ZIC1* driver gene^35^, which is also a potential drug target^50^. It is known that ASM plays an important role in the regulation of allele specific expression (ASE)^51^ and ASM is increased in cancer, caused by disease associated regulatory SNPs^52^. A total of 20 genes show ASM as well as significant ASE effects (FDR < 0.1, methods), where increased methylation is associated with reduced expression (Pearson R: −0.471, p-value: 3.6 × 10^−2^, **Figure 3E**), when accounting for haplotype copy number state this correlation is stronger (Partial correlation R: −0.501, p-value: 2.8 × 10^−2^), again we observe a significant overlap between ASE and ASM genes (Fisher’s exact test statistic: 4.1, p-value: 2.63 × 10^−6^, using all genes expressed in primary tumor as background).

### Haplotype resolved functional interpretation of complex rearrangements

We notably observed ASM also in association with the chromothripsis event resulting in the complex CS11-17 structural segment. Since the CS11-17 rearrangement occurs in only one haplotype, we searched for ASM between the CS11-17 haplotype and the corresponding wild-type (non-rearranged) haplotype stretches. We find a global pattern of demethylation of the CS11-17 haplotype in contig 2 (**Figure 4A**) compared to the non-rearranged haplotype, which includes demethylation of *TRIM66* and *STK33*. On contig 1 of CS11-17, the promoter regions of *SPATA32, USP22* and *MAP3K14-AS1* are demethylated on the corresponding wild-type haplotype in the primary tumor, while being methylated on CS11-17 as well as on both of the unaffected haplotypes in the relapse (**Figure 4B**). No ASE is found for the genes on the demethylated contig 2 of CS11-17. *USP22* on contig 1 of CS11-17 shows higher ASE in the demethylated allele, and *MAP3K14-AS1* in the methylated allele, most likely driven by the higher copy number of the chromothriptic haplotype.

**Figure 4.**
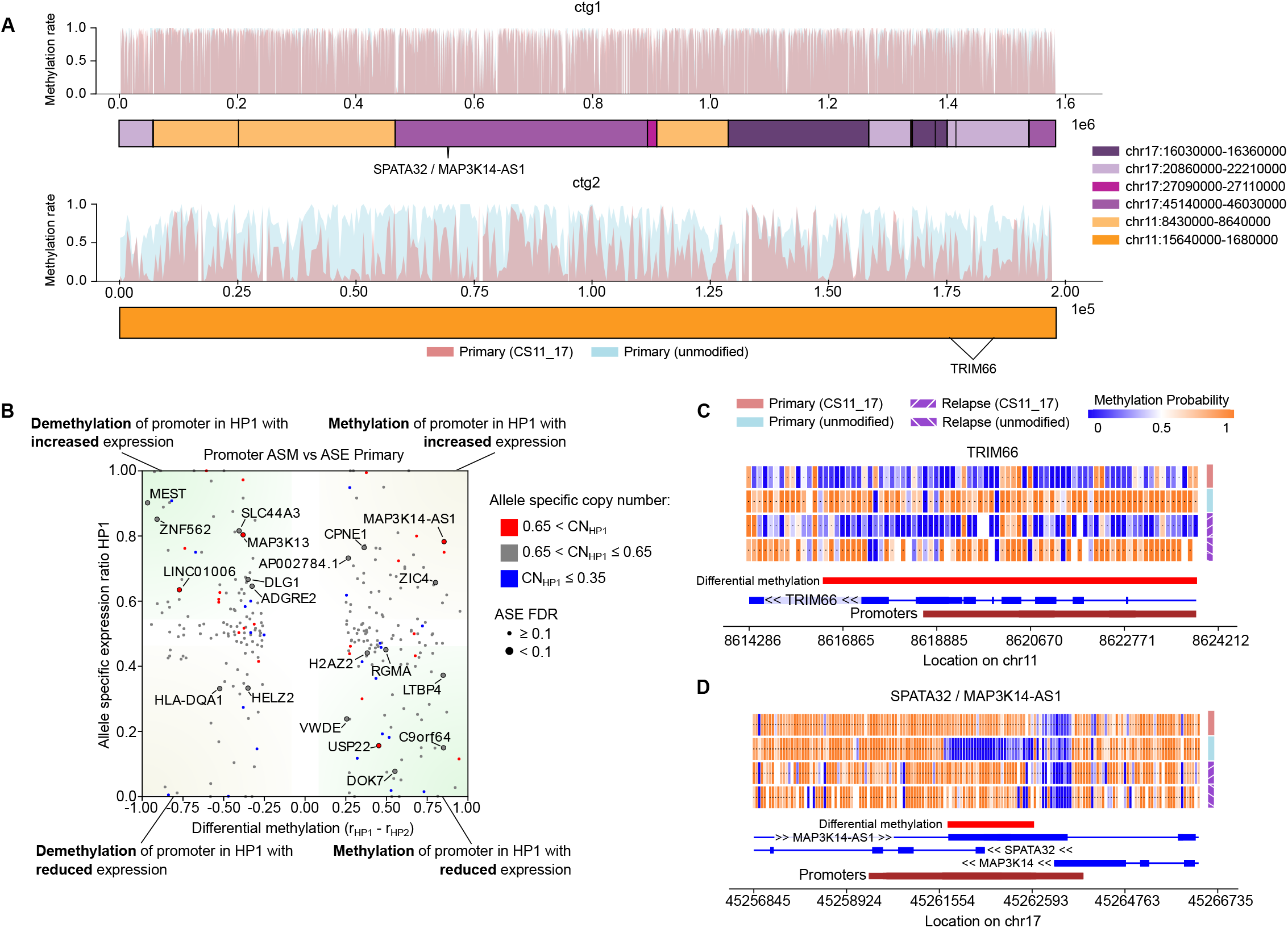
Methylation of complex genomic rearrangements. **(A)** Methylation rates of chromothriptic contig CS11_17 in the primary tumor sample show global demethylation of contig2, containing genes TRIM66 and STK33, to a methylation rate of 42% on the CS11_17 haplotype from 76% in the corresponding genomic ranges on the non-chromothriptic haplotype. While contig1 displays some allele specific differences, no significant global effects are detected. **(B)** ASE and promoter linked ASM in primary tumor. **(C)** Demethylation of CS11_17 haplotype of contig2 effect shown on TRIM66 promoter. **(D)** ASM of promoter of gene SPATA32 and antisense transcript MAP3K14-AS1 on contig1.

### Functional annotation of the templated insertion threads and telomere SVs

We next performed similar functional annotation of the templated insertion threads and the telomere insertions. The templated insertion threads appear to retain their original methylation state with only a slight reduction in methylation rate measured (average methylation rate reduction structure 1: 0.16, structure 2: 0.09, **Figure S17**). Interestingly the first templated insertion thread (**Figure 2B**) lands in an intronic region of *BASP1*, which was previously implicated in metastatic medulloblastoma in a mouse model specifically by transposon insertion mutagenesis^53^. While this is a different type of insertion, we notably do observe differences in splicing of *BASP1* between the samples. Within the relapse sample, which does not harbor the templated insertion thread, we observe three splice junctions that are not used in the primary tumor (Junction 1 (5:17260615-17275208): Fisher’s exact test p-value: 1.5 × 10^−23^, Junction 2 (5:17228332-17275208): p-value: 2.0 × 10^−22^, Junction 3 (5:17263478-17275208): p-value: 4.4 × 10^−10^). The junction used for the main *BASP1* isoform (*BASP-201*) is more frequently used in the primary tumor as compared to the relapse (**Table S9**). To further explore the functional relevance of the observed templated insertion threads we also searched for potential gene dysregulation effects within the transcriptomic data available for liposarcoma samples in PCAWG^2^. We identified one liposarcoma sample (donor id DO219945), which harbors a templated insertion thread on chromosome 12 whose breakpoints intersect the coding sequence of proliferation-associated protein 2G4 (*PA2G4*), which can act as a contextual tumor suppressor^54^, in association with reduced *PA2G4* expression (**Figure S18A**). Another liposarcoma sample (donor id DO219967) shows strong overexpression of *CCND3*, a known sarcoma oncogene, and *BYSL*, a gene associated with tumor prognosis^55^, in the immediate vicinity of a templated insertion thread (**Figure S18B**). These examples suggest a possibly relevant role of template insertion threads in cancer, illustrating the need of routinely generated long reads to fully characterize somatic SVs with respect to cancer-related genes in tumor genomes.

Analyzing the telomere-associated SVs we find that four of such SVs observed in the primary tumor and relapse samples (**Figure 2G, S13**) harbor a breakpoint junction in intronic regions of protein coding genes, namely *TLL1, THADA*, and *MYPOP* in the primary tumor and *LUZP2* in the relapse sample. The *MYPOP* and *TLL1* SVs also show short templated insertions between the telomeric part and the above mentioned genes, with templated insertion source sequences originating from intronic regions of various other genes (**Figure 2G, S13**). We performed differential expression analysis between the primary tumor and relapse, and found that *TLL1* showed a slightly reduced expression in the primary tumor (−1.15 l2fc) whereas *LUZP2* and *MYPOP* displayed a reduced expression in relapse (−1.16 l2fc and −1.08 l2fc, respectively). Additionally, *MYPOP* is found to be subclonally amplified in the haplotype where the telomere associated SV is observed (allele specific copy-number ratio 0.7) with a matching allele specific expression rate (0.75). This amplification extends across most of chromosome 19q and happens only in the primary tumor, while in relapse the copy number ratio for *MYPOP* is 0.53 (**Figure S21)**.

## Discussion

We describe the haplotype-resolved genetic and epigenetic profile of a diagnosis and post-therapy medulloblastoma using long reads and present new computational methods for targeted *de novo* assembly and complex SV characterization, as well as phasing, segmentation, and investigation of ONT methylome profiles. We used an integrated phasing approach that combines long-reads with statistical phasing for haplotyping which enabled the assembly of a 1.55 Mbp chromothripsis event spanning 14 breakpoints. Furthermore, by leveraging the joint genetic and epigenetic readout of ONT data, we revealed haplotype-specific and chromothripsis related methylation changes – analyses difficult to pursue with short reads due to the sparsity of germline heterozygous singlenucleotide polymorphisms and limitations in read length. The combination of long read genetic and phased methylation information from ONT reads can further be used to detect aberrant expression patterns, such as allelic expression imbalance or gene fusion events at greater level of detail. In the future, deep coverage and highly accurate long-read data will be needed to achieve the complete *de novo* assembly of cancer genomes, especially in the context of intra-tumour heterogeneity, contamination of normal cells, and large numbers of complex rearrangements.

The proposed long-read methods enabled us to describe a new complex DNA rearrangement pattern, termed templated insertion thread, consisting predominantly of short segments (<1kbp) that are copied and (self-)concatenated into amplified, highly repetitive somatic sequences of up to 50kbp in size. Umbreit et al. did not detect self-concatenating insertions of high copy-number in the cell cultures of their *in vitro* study, and their recently described tandem short template jump pattern^22^ therefore bears differences to the template insertion thread pattern described here. However, the study by Umbreit et al.^22^ provided additional validation data from a renal cell carcinoma, which included an example of a chained rearrangement with a zig-zag pattern of templated insertions involving at least a few self-concatenations. These validation data, therefore, further support the templated insertion thread pattern defined in our study. Future analysis of larger sample sets using long-reads will be required to delineate the full extent and scope of concatenated insertions in cancers, which is likely to be currently underestimated. Notably, tandem short template jumps^22^, like templated insertion threads, show an association with chromothripsis – which leaves the possibility of a continuum of concatenated insertion patterns arising in conjunction with complex DNA rearrangement processes.

We demonstrate using a new graph-based method, *rayas*, that templated insertion threads can be identified in short read WGS data, which is important as it allows further study of this complex rearrangement pattern in existing large short-read cancer genomic cohorts. We describe a remarkable enrichment of this pattern in different adult cancers, with the strongest prevalence in liposarcomas (74% of cancer samples affected) and a clear colocalization of these events with genomic regions undergoing giant marker chromosome formation and chromothripsis. We did not identify any additional medulloblastoma samples with templated insertion threads in the PCAWG short read dataset, which is perhaps explained by the relatively low portion of medulloblastoma samples contained in the PCAWG cohort exhibiting chromothripsis (~12%)^56^. One note of caution is that discovery of high-complexity regions as seen in templated insertion threads using shortreads is obscured by somatic SV calling pipelines because multiple distinct SVs co-occur at the same SV breakpoint leading to algorithmic clustering and SV merging issues. This is contrary to long reads that have the capability to fully resolve the complex structure and composition of structural rearrangements in cancer genomes. While *rayas* can overcome this issue in part, it is likely that short read WGS masks additional cases of templated insertion threads, especially where they involve short (<1kb) templated insertion units or repeat-rich DNA, given the relatively poor sensitivity of Illumina reads for calling such SVs^13^.

The long-read data also enabled investigation of the association of complex SVs and telomeric repeats, an analysis that revealed the fusion of telomeres with chromosomes that underwent chromothripsis. Some of these events were captured in a single long ONT read connecting a telomere to various SV rearrangements, reminiscent of SV mutations stabilized by independent telomere fusions. The assignment of telomeric repeats to chromosomal haplotypes also highlighted the need for continuous reference improvements, as some of these events could only be unambiguously resolved using the new CHM13 telomere-to-telomere (T2T) assembly^31^. A comparable analysis on short-read data failed to resolve the telomere-associated complex rearrangements, and only three out of the five SV to telomere junctions showed confident telomeric repeat motifs in an unmapped mate or a soft-clipped read, which underscores the critical need for long-read sequencing to investigate telomere-associated structural rearrangements, which are considered a key cancer mutational process in association with telomere crisis^28^.

Despite the unprecedented view into somatic SV rearrangement patterns that ONT long-reads enable, a few key challenges remain: 1) Our strategy focused on targeted assemblies of high-copy number regions due to the moderate long-read sequencing coverage (up to 30-fold); while long-read sequencing remains costly compared to Illumina sequencing, future gains in throughput will enable studies in larger sample panels with coverages adequate for uncovering SVs in the context of intra-tumor heterogeneity. 2) Our assemblies failed to resolve peri-centromeric regions involved in the CS11-17 chromothripsis region exceeding the available read length. As ONT read lengths are determined by the sample preparation protocol, this suggests that “ultra-long” preparations may prove beneficial to characterize somatic SVs contained within repeat-rich regions, once available for routine application. 3) Further computational methods development will be needed to achieve the assembly of entire derivative chromosomes in cancer, including new algorithms for SV-aware haplotyping and multi-allelic assemblies.

In summary, our study shows the benefits of using long reads in refining complex and repetitive rearrangement patterns such as templated insertion threads and telomere associated SVs, and to integrate these with allele-specific methylation and expression changes. The computational methods developed in our study provide the foundation for a more broad application of long reads in cancer genomics to uncover new somatic mutation patterns, and pave the way for deciphering the complex relationship of genetic and epigenetic changes in cancer biology.

## Data Availability

Sequence data have been deposited at the European Genome-phenome Archive under the accession number EGAS00001005410.

## Software Availability

Lorax: https://github.com/tobiasrausch/lorax

Rayas: https://github.com/tobiasrausch/rayas

Wally: https://github.com/tobiasrausch/wally

Analysis scripts: https://github.com/PMBio/mb-nanopore-2022/

## Acknowledgements

We thank Frauke Devens, Kim Judge as well as DKFZ and EMBL IT and sequencing core facilities for excellent technical support. The present contribution is supported by the Helmholtz Association under the joint research school “HIDSS4Health - Helmholtz Information and Data Science School for Health”.

A.E. received funding from the DFG (project number 460595631) and from the Wilhelm Sander Foundation (project number 2020.115.1). J.O.K. received funding from the BMBF (031L0184C) and from the NIH (1R01HG010169-01 and 2U24HG007497-05).

## Author contributions

E.B., O.S., A.E. and J.O.K. designed the study. A.L. performed long read base calling and alignment. R.S. performed methylation calling and differential methylation analysis, T.R. implemented phasing, targeted assembly workflows, germline and somatic variant discovery and complex structural variant calling. R.S. and M.J.B. performed RNA alignment and expression quantification and performed subsequent expression analyses. M.S. performed FISH and established xenograft models for metaphase spreads. T.R., R.S., M.J.B., A.E. and J.O.K. analyzed complex mutation patterns and targeted assemblies. R.S. implemented the gene fusion validation. T.R. and J.O.K. performed templated insertion analysis and interpretation in PCAWG. R.S., T.R. and M.J.B. prepared the main display items, with additional contributions from A.E. and J.O.K. T.R., R.S., M.J.B., A.E. and J.O.K. wrote the manuscript, with input from E.B., A.L. and O.S.

## Declaration of interests

E.B. is a paid consultant and shareholder of Oxford Nanopore Technologies (O.N.T.). A.L. has received financial support from O.N.T. for consumables during the course of the project and is currently an employee of Oxford Nanopore Technologies (O.N.T.). The remaining authors declare no competing interests.

## Methods

### Patient material, DNA extraction and short-read whole-genome sequencing

All biological samples included in this study were obtained after receiving written informed consent in accordance with the Declaration of Helsinki and approval from the respective institutional review boards. Medulloblastoma samples used for bulk sequencing had a tumor cell content confirmed by neuropathological evaluation of the hematoxylin and eosin stainings. DNA was extracted from frozen tissue and from blood using Qiagen kits. Purified DNA was quantified using the Qubit Broad Range double-stranded DNA assay (Life Technologies, Carlsbad, CA, USA). Genomic DNA was sheared using an S2 Ultrasonicator (Covaris, Woburn, MA, USA). Short-read whole-genome sequencing and library preparations for tumors and matched germline control were performed according to the manufacturer’s instructions (Illumina, San Diego, CA, USA). The quality of the libraries was assessed using a Bioanalyzer (Agilent, Stockport, UK). Sequencing was performed using the Illumina X Ten platform.

### DNA methylation array data

Medulloblastoma samples were analyzed using Illumina Infinium HumanMethylation450 BeadChip (450k) arrays or Methylation BeadChip (EPIC) arrays according to the manufacturer’s instructions.

### RNA sequencing

RNA was extracted from frozen tissue using Qiagen kits. RNA quality was assessed using a Bioanalyzer (Agilent, Stockport, UK). Short-read RNA sequencing and library preparations for tumors were performed according to the manufacturer’s instructions (Illumina, San Diego, CA, USA). The quality of the libraries was assessed using a Bioanalyzer (Agilent, Stockport, UK). Sequencing was performed using the Illumina platform.

### Fluorescence in situ hybridization (FISH)

Nick translation was carried out for BAC clone RP11 651L9 (chromosome 17) and centromere 17. FISH was performed on metaphase spreads from patient-derived xenograft models or tumor tissue using fluorescein isothiocyanate-labeled probes and rhodamine-labeled probes. Pre-treatment of slides, hybridization, post-hybridization processing and signal detection were performed as described previously^57^. Samples showing sufficient FISH efficiency (>90% nuclei with signals) were evaluated. Signals were scored in, at least, 100 non-overlapping metaphases or nuclei. Metaphase FISH for verifying clone-mapping position was performed using peripheral blood cell cultures of healthy donors as outlined previously^57^.

### Long-read sequencing

DNA was quantified using Qubit (Thermo Fisher) and fragment size assessed using FEMTOPulse (Agilent). Libraries were prepared using SQK LSK-109 (Oxford Nanopore) following the manufacturer’s protocol and sequenced on the PromethION (Oxford Nanopore).

### Short-read alignment, variant calling and copy-number segmentation

Paired-end, short-read FASTQ files (2×151bp) were aligned to the GRCh38 reference genome using the alternate contig-aware bwakit^58^. Alignments were sorted and indexed using samtool^s59^ and quality-controlled with alfred^60^. The median coverage of the blood (control), primary tumor and relapse sample were 48x, 45x and 47x, respectively. The insert size ranged from 373bp to 406bp for the three samples.

Single-nucleotide variants (SNVs) and short insertions and deletions (InDels) were called using FreeBayes^61^ and Strelka2^62^. For germline variants we used a consensus approach and only retained polymorphisms supported by FreeBayes and Strelka for subsequent haplotyping. The integration of these two short-read germline call sets on GRCh38 yielded 3,790,471 bi-allelic SNVs and 568,168 bi-allelic insertion and deletions. Bcftools was used to normalize and left-align indels. Copy-number segmentation employed Delly’s cnv mode^63^ with the GRCh38 mappability map and the DNAcopy^64^ package of the Bioconductor project (**Figure S3**). Structural variants were called using delly^63^ in a paired tumor-normal fashion to distinguish germline and somatic SVs. All command-line tools were installed using bioconda^65^.

### Long-read alignment and variant calling

Long reads from Nanopore sequencing were basecalled with guppy version 4.0.14 using the high accuracy model for PromethION (r9.4.1_450bps_hac_prom). Resulting FASTQ files were aligned to the human reference genome (GRCh38) using minimap2^66^ using the ‘--ax map-ont’ option and otherwise default parameters. The median long-read coverage was 15x for the blood and relapse sample and 30x for the primary tumor. The median read length was 4,480bp, 4,993bp and 5,678bp for the blood, primary tumor and relapse sample, respectively. The estimated sequencing error rate of the aligned data using Alfred’s qc mode^60^ was estimated to be 8.4% for the blood sample and 6.8%-6.9% for the tumor samples.

Structural variants (SVs) from the long-read data were called using Nanovar^67^, Sniffles^68^ and Delly^63^. Consensus germline SVs were filtered using a stringent reciprocal overlap of 80% and a maximum breakpoint offset of 50bp, yielding 7,952 deletions and 8,185 insertions, which is lower compared to recent studies using long-reads^12,13^ likely because of our relatively low germline coverage of only 15x (**Figure S1**). For somatic SVs we followed a more lenient union approach of short-read SV calls (delly) and long-read SV calls (nanovar, sniffles and delly) to not miss any interesting variants and only required absence of an SV in the matched control and a minimum support of 2 reads in the tumor, followed by manual inspection of somatic SVs in IGV^69^ and a newly developed alignment visualization tool, called *wally*, which enables a fast batch alignment plotting of hundreds of SVs in a paired tumor-normal split-view.

### Nanopore methylation calling

Read-level CpG methylation likelihood ratios were estimated using nanopolish^70^ version 0.11.1. Methylation rates were computed from binarized methylation calls thresholded at absolute loglikelihood ratio of 2.5 and compared to methylation rates observed in 450k arrays. Methylation ratios predicted from long reads showed good correlation with array data, with pearson R 0.9453 for the primary tumor sample and R 0.9141 for the relapse sample.

### Haplotype-phasing of short variants

We used a three-stage approach to phase bi-allelic heterozygous SNVs and InDels present in our consensus call set from FreeBayes and Strelka. In brief, the first stage uses read-based phasing of the long-read data to generate initial haplotype blocks, these are concatenated using population phasing in the second step and finally, remaining switch errors are corrected using shifted allelic ratios in the matched tumor. The procedure is illustrated in **Figure S2** where initial phased blocks are colored red and blue that are then extended using statistical phasing and corrected based on the matched tumor genome.

For read-based phasing we used WhatsHap^71^ with the ‘--indel’ option and the aligned long-read data. The WhatsHap output VCF was indexed using HTSlib^72^. WhatsHap determines phased sets which are groups of heterozygous genotypes at which the phase has been inferred using long reads. These phased sets are specified in the PS field of the VCF/BCF file format^73^. With the SHAPEIT4 algorithm^74^ and the phased blocks from WhatsHap we then carried out population phasing using the 1000 Genomes haplotype reference panel^20,75^. We used the ‘--use-PS 0.0001’ option to define the expected error rate in the phased sets. The statistically phased VCF files were then augmented for each variant with the matched tumor B-allele frequencies to correct remaining switch errors in regions of unequal haplotype ratio in the tumor sample. As a result of statistical phasing and the use of a haplotype reference panel the statistically phased VCF files are restricted to high-quality variants present in the panel. We therefore used this phased VCF file as a haplotype scaffold to drop in additional variants present in our donor using WhatsHap and the long-read aligned data. Overall, our haplotype-phasing approach phased 2,642,137 bi-allelic heterozygous variants (2,214,532 SNVs and 360,226 InDels) at a median read length of approximately 5kbp which allowed us to study almost the entire mappable genome, 93.59% for the primary tumor and 90.89% for the relapse, in a haplotype-resolved manner. To split alignment files by haplotype we employed Alfred^60^ using the phased VCF and the unphased alignment as input.

### Targeted assembly of complex DNA rearrangements

To enable targeted assembly of complex SVs, we used our haplotype scaffold and the integrated map of somatic structural variants and copy-number alterations. We first applied delly’s cnv mode and the somatic SV calls to identify amplicons on chromosome 11 and chromosome 17 that are inter-connected by split-reads and that have approximately the same total copy-number. We then developed a targeted method to assemble these high copy-number regions by selecting reads that either bridge at least two amplicons or are part of the amplified haplotype based on the depth observed for each germline allele. We implemented the method in our long-read analysis toolbox for cancer genomics, termed *lorax*, and the tool requires as input the phased germline variants in VCF/BCF format, a set of amplicon regions in BED format and the input tumor BAM file. The method then screens the BAM file for split-reads connecting at least two amplicons and it annotates the haplotype support based on all phased, heterozygous variants covered by the read sequence. Each read is then assigned to either haplotype 1 or haplotype 2 based on the observed variants. The total allelic depth across all reads in the respective amplicon region determines the amplified haplotype which is retained for further analysis. We discard all reads that have confident alignments outside the amplicon boundaries to deplete reads from contaminating normal cells occurring on the same haplotype background or sub-clonal reads from different rearrangement structures. User-defined parameters control the precision of amplicon boundaries (default 1kbp), the minimum required clipping length of split reads (default 100bp) and the minimum mapping quality (default 10). A final pass through the BAM file extracts the sequences of all selected reads, which are then assembled and polished using wtdbg2^76^. *Lorax* also re-estimates the amplicon boundaries based on the observed read clipping patterns which was used to iteratively refine the input amplicon regions. We trimmed the assembly at repetitive ends that lacked a unique alignment to the reference. The final contigs were aligned back to the reference genome using minimap2^66^ to infer alignment coordinates and breakpoints.

### Discovery of complex templated DNA rearrangements

To discover complex templated DNA rearrangements using short-reads we devised a graph-based algorithm, called *rayas*, that uses matched tumor-normal cancer genomics sequencing data. The algorithm parses the tumor and normal BAM file to compute a sample-specific coverage and splitread profile at single-nucleotide resolution. Rayas uses soft- and hard-clips and records the positions where these splits occur. The coverage profile is used to determine the average genomewide coverage, its standard deviation and to normalize for overall coverage differences between tumor and normal. Using a minimum seed window size (default 100bp) rayas then scans the coverage profile for putative SV breakpoints, always screening two adjacent windows for unexpected coverage increases when entering a templated insertion source segment or unexpected coverage decreases when leaving a templated insertion source segment. Command-line parameters control the minimum number of split-reads required at these SV breakpoints and the required magnitude of the coverage increase or decrease. The matched control is processed simultaneously to account for potential mapping artifacts, i.e. regions where both the tumor and the control show unexpected coverage and split-read patterns which are subsequently filtered out. Once all candidate segments have been identified, rayas re-uses the identified split-reads to connect segments and builds a graph *G* = (*V, E*) with *v ϵ V* representing a templated insertion source segment and *e* = (*v, w*) *ϵ E* being an edge from *v* to *w* with *weight*(*e*) representing the split-read support. Using the connected components of *G, rayas* filters out singletons (i.e. segments lacking confident split-read support) as well as connected subgraphs *G_s_* = (*V_s_, E_s_*) with *V_s_* ⊆ *V* and *E_s_* ⊆ *E* where all nodes of *G_s_* are nearby in the genome with the definition of nearby depending on a user-defined threshold (by default 10kbp). All remaining connected components are written to a BED file with a unique component id. For each component, all genomic segments and edges are outputted and the results can be visualized as a graph (**Figure S11**). Using this approach we identified two templated insertion threads in the primary tumor. In addition, a single additional putative instance of this pattern was detected in the Illumina data of the relapse but not in the ONT data from the same sample; this putative event showed much lower split-read support (5 compared to >>100 for the primary tumor templated insertion threads) and an unexpected density of variant calls, suggesting that it may be caused by a mapping artifact or a collapsed repeat rather than a templated insertion thread. A simple threshold for the minimum split-read support (i.e., node out-degree in the rearrangement graph) removes such false positives, indicating excellent sensitivity and specificity of *rayas* using illumina data. For the PCAWG data, we filtered for connected components with at least one segment with a total copy-number greater than 10, a node degree greater than 50 and evidence of at least one direct self-concatenation supported by at least 3 split-reads, as these features were characteristic of the templated insertion threads found in the medulloblastoma.

The algorithm implemented in *lorax* for detecting templated insertions with long reads uses the same discovery approach as *rayas*, but then scans the original alignment data to extract long reads that span multiple templated insertions. These reads can be selectively assembled, inspected through self-alignments or back-aligned to the source sequence segments as shown in **Figure 2**. The visualization of long read alignments spanning dozens to hundreds of breakpoint junctions employed minimap2^66^, MUMmer^77^, custom R scripts and a newly developed tool, called *wally*, that enables the plotting of long read mappings with alignments widely distributed across the genome by lining up matches along the read sequence (as shown in **Figure S9**).

### Telomere analysis of derivative chromosomal segments

As part of our long-read analysis toolbox for cancer genomics, termed *lorax*, we also developed a method that identifies telomeric motifs, repeats of TTAGGG, TGAGGG, TCAGGG, TTGGGG or their reverse complement, in error-prone ONT reads and applied this method to the long read data of the primary tumor and the relapse sample. As suggested previously^78^, we start by precomputing all possible strand-specific 18-mer telomere motifs, scan all long-reads for exact motif matches and count their occurence. We then search for distal non-telomeric alignments of these reads and intersect reads that show both a telomeric repeat and a unique alignment outside a telomere region of a minimum length of 1kbp. We use the control genome to filter out likely mapping artifacts due to incomplete reference sequences by masking alignments from the control genome that show both a telomeric repeat and a unique alignment outside a telomere region. In case of mapping ambiguities, we used the CHM13 telomere-to-telomere (CHM T2T) assembly ^31^ as an alternative reference sequence. The method to detect telomere fusions is implemented in our long-read alignment toolkit *lorax* as a new sub-command. For the matched illumina data, we apply a window-based search (default 1kbp) that counts reads with a telomeric motif based on the mapping location of the read (or its mate if the read is unmapped). If both read1 and read2 are unmapped the sequencing pair is discarded. We filter out all windows that are discovered in the matched control (blood) and retain in the tumor only windows with at least 5 supporting paired-ends. The short-read method is implemented in the alfred toolkit^60^ as a new sub-command, called ‘alfred telmotif.

### Differential methylation testing

In order to find genomic regions with differential methylation between samples, we used the software package PycoMeth^34^. PycoMeth aggregates methylation likelihood ratios reported by Nanopolish over predefined regions, computes a read-level methylation rate from thresholded loglikelihood ratios (threshold 2.0) and then performs a Wilcoxon rank-sum test (for 2-sample comparison) or Kruskal Wallis test (for more than two samples) for methylation rates across samples. P-values were then adjusted for multiple testing using independent hypothesis weighting^79^, using a weight based on the variance of methylation rates, and the Benjamini-Hochberg method^80^. Regions with FDR<=0.05 are reported as differentially methylated regions (DMRs). Candidate regions for differential methylation testing are selected based on two different segmentation methods: 1) sequence segmentation and 2) methylome segmentation. Sequence segmentation uses PycoMeth’s CGI Finder module, which determines CpG islands based on local CG-density. For methylome segmentation PycoMeth Meth_Seg, a *de novo* methylome segmentation method which implements a bayesian changepoint-detection algorithm, is used to determine regions with consistent methylation rate from the read-level methylation predictions. For ASM analysis, PycoMeth Meth_Seg was provided with haplotype information to perform a haplotype-aware segmentation.

We investigated differentially methylated regions between the primary tumor and the relapse sample, as well as between all three samples by applying PycoMeth Meth_Comp using both candidate region approaches with the parameter using the parameter `—hypothesis bs_diff` in order to test for difference in read-level methylation rate per segment. DMR identification was performed both in a sample and haplotype comparison mode. To assign reads to haplotypes we used WhatsHap’s `haplotag` command and the three-stage phased blood variants. This haplotype assignment was used as the read-group parameter in PycoMeth, allowing it to consider ASM in the methylome segmentation. In PycoMeth, differential methylation calling was then performed between haplotypes within each sample, in order to determine regions with ASM. For further analyses, DMRs were filtered by an effect size threshold of 0.5 absolute methylation rate difference. Differentially methylated regions were then mapped to genes based on their proximity to a transcription start site (TSS), that is they were labeled as promoter methylation if a region was in the range 2,000bps upstream to 500bps downstream from the any transcript’s TSS, or if it overlapped with an enhancer active in Cerebellum as annotated by EnhancerAtlas 2.0^81^. Enhancers were then linked to the nearest gene, if the gene is closer than 30kbps. Since detection power in the relapse sample was lower, due to lower read-depth, we investigated whether ASM effects found in primary tumor could be found in relapse as well by applying the same 0.5 absolute methylation rate difference threshold.

### RNA alignment and expression quantification

Gene-expression quantification was performed in line with the GTEx standards. In short, we (re)processed the RAW expression data by first aligning the reads to the human reference genome, build 38, using STAR in two step mapping per sample. The mapping was performed in two modes. One for the allele specific expression, using a custom reference genome, replacing the homozygous SNP variants with the relevant genotype of the sample, and supplying a VCF with heterozygous variants when mapping in STAR, used for allele specific expression and gene fusion detection. Second, for the differential expression and splicing analyses we remapped the samples to the standard genome. Gene information was taken from ENSEMBL (v101) and gene-expression quantification was performed using RNASeQC, in line with the GTEx consortium expression quantification. Using LeafCutter^82^ we quantified splicing across the two samples, as well as a cerebellum reference dataset (SRP151960)^83^.

### Reference RNA expression datasets and differential expression

For comparative expression analysis we leverage data from the ALS consortium (SRP151960^)83^ and GTEx cerebellum expression data. The data from the ALS consortium were reprocessed as done for the two medulloblastoma samples, see above, and the GTEx data^84^ was used as is. This data was leveraged both for direct comparison of expression levels, and for correction of the gene expression levels.

The first five principal components (PCs) were calculated on the combined ALS and GTEx dataset. The medulloblastoma samples were projected into this same PC space, using the rotation information, and the first five PCs were regressed out from the expression levels of all samples, medulloblastoma, GTEx and ALS. Next we used a Z-score transformation on both the raw and corrected expression of the reference samples and placed the two medulloblastoma samples in these distributions. Given that there are still major differences between the samples and studies, in terms of age, disease and batch, we only use the two samples in a comparative setting. The reference data is used to test for concordance of effects with and without correction. For the differential expression analysis we used the log TPM values and checked concordance in Z-scores.

### Allele specific expression and allele specific copy number estimation

ASE on the primary tumor and relapse samples was called from the RNA sequencing data using WASP^85^ and the phased germline variants, using the approach described in the WASP paper^85^. In order to verify whether ASE was driven by DNA copy number amplification or depletion in one haplotype, we estimate allele specific DNA copy number ratio using GATK CollectAllelicCounts^86^ on the same variants used to identify ASE.

### Gene fusion and validation using DNA long reads

Potential gene fusions were detected from RNA sequencing data using Arriba^87^ (V2.0.0). The SVs called from both short and long read data were used to inform Arriba, and we included the provided blacklist, other settings were left at defaults. After identification of the gene fusion pairs we set-out to validate these using the long read DNA data. First, we check for individual read support from ONT reads with chimeric alignments mapping to both genes. Fusion pairs involving long intergenic non-coding RNA genes, which are characterized by long introns of on average 10kbps length^88^, or fusion containing large intronic insertions, however often do not have individual genomic reads spanning exons of both genes. In order to additionally validate such fusions with large insertions, for which no single ONT read spans the fusion pairs, we devised a graph-based method to suggest the most plausible gene fusion reconstruction. We construct a graph with nodes representing each base pair position in the reference and edges representing neighboring basepairs. Structural variations, both inter- and intrachromosomal, were then represented as additional edges in the graph, creating shortcuts between the locations on the side of the genomic breakpoint connected by the structural variation). A gene fusion pair was then explained by determining the shortest path between the two fusion partners in the graph using Dijkstra’s algorithm for shortest paths^89^. Edges which crossed the exons of a gene not involved in the fusion were removed for the purpose of finding the shortest path. Fusion pairs were classified as either validated by individual read support, explainable using the graph algorithm, or both (high confidence read support).

## Notes

https://ega-archive.org/studies/EGAS00001005410

