## Supplementary Figures and Supplementary Tables S1-S5 for "Long-read sequencing of diagnosis and post-therapy medulloblastoma reveals complex rearrangement patterns and epigenetic signatures"

### Supplementary Materials

Supplementary materials for the manuscript “Long-read sequencing of diagnosis and post-therapy medulloblastoma reveals complex rearrangement patterns and epigenetic signature”.

This supplementary information is structured in figures and tables.

**Supplementary Figures**


S1. Germline structural variant size distribution

S2. Haplotype-phasing approach

S3. Copy-number profiles

S4. Copy-number plots of chromothripsis chromosomes

S5. FISH analysis

S6. Self-alignment of the templated insertion thread

S7. Templated insertion thread alignments

S8. Subsampling analysis of ONT reads spanning templated insertion threads

S9. Genomic matches of a single ONT read with a templated insertion thread

S10. Tumor heterogeneity of templated insertion threads

S11. Templated insertion rearrangement graphs

S12. Templated insertion threads across 2,569 cancer genomes

S13. Telomere sequences associated with rearranged genomic regions

S14. Comparing 450k array versus Nanopore methylation calls

S15. Length of differentially methylated regions

S16. Methylation array coverage of differentially methylated regions

S17. Methylation of templated insertion threads

S18. Gene expression effects of templated insertion threads

S19. Differential promoter methylation compared to differential expression

S20. Differential methylation of TBX1

S21. Allele copy number ratios for chromosome 19 with telomere associated SV

**Supplementary Tables**

S1. Long-read sequencing statistics

S2. Short-read sequencing statistics

S3. FISH analysis using probes for RP11-651L9 and centromere 17

S4. Combined FISH analysis using probes RP11-651L9 and centromere 17

S5. FISH on metaphase spreads from matched patient derived xenographs

S6. Templated insertion threads identified in PCAWG

S7. Overview of expression effects, methylation effects, and genetic variants between samples and between haplotypes

S8. Allele specific expression, allele specific methylation, and allele specific genomic copy number of primary tumor sample

S9. Overview of the leafcutter analysis on BASP1

### Supplementary Figures

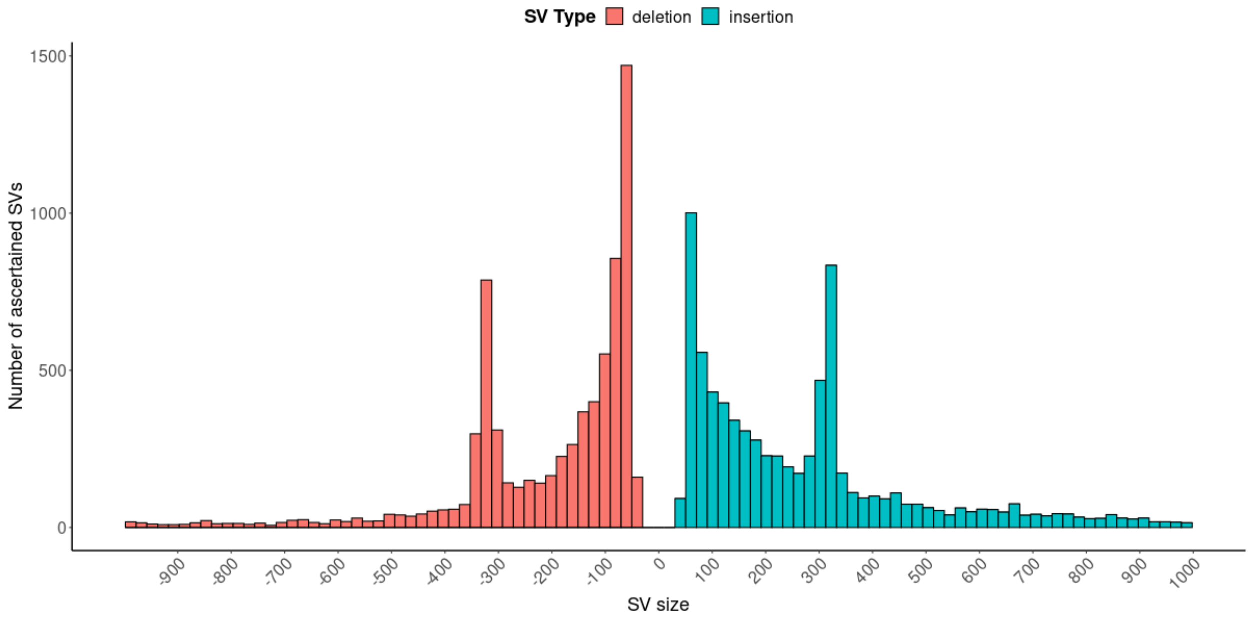


**Figure S1. Germline structural variant size distribution.** Germline structural variants >50bp binned by size and type. Deletions are shown to the left (negative size) and insertions to the right (positive size). Insertions and deletions show the characteristic ALU peak at approximately 300bp, indicative of transposable element insertions and deletions relative to the reference.

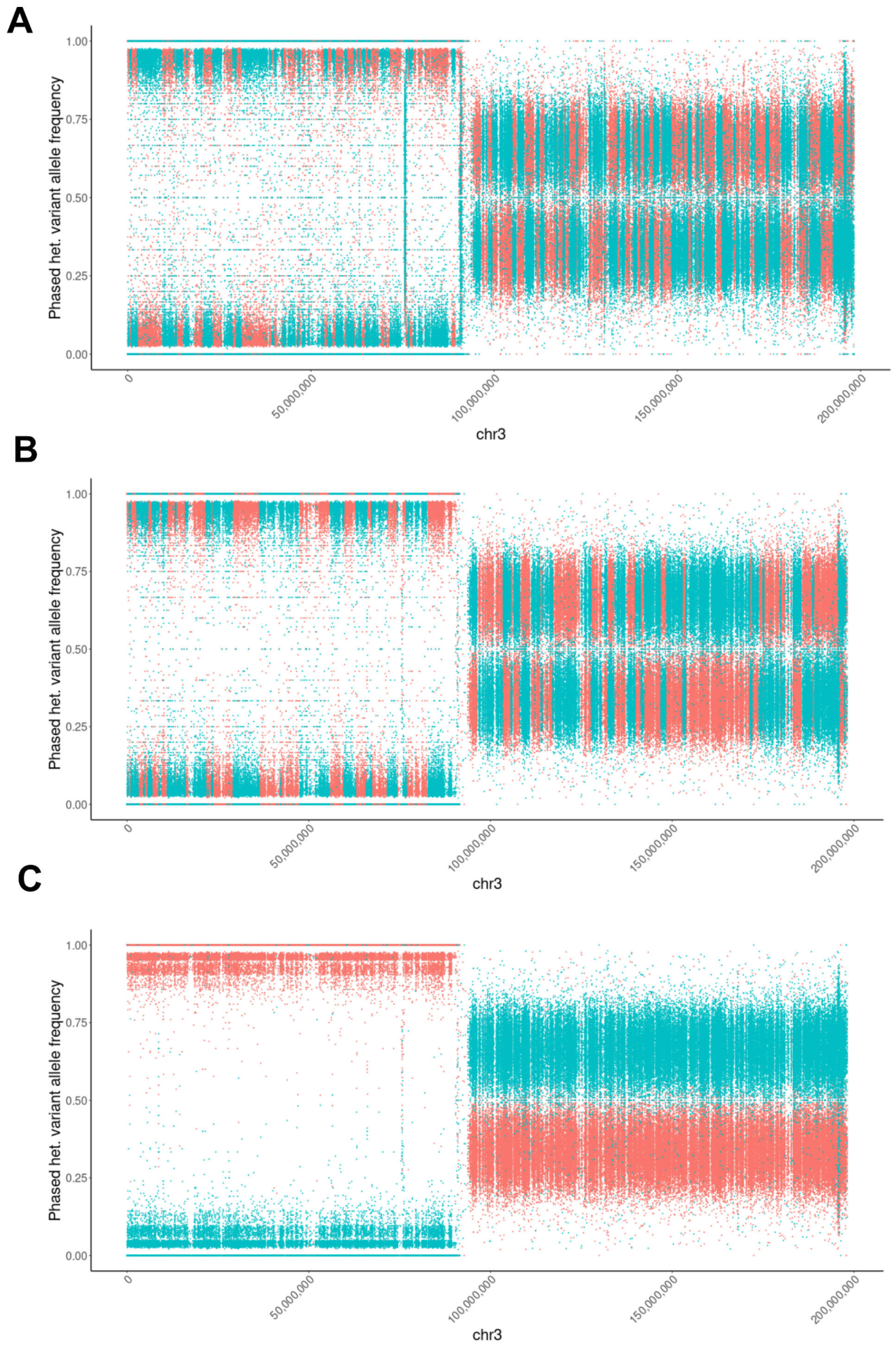


**Figure S2. Haplotype phasing approach.** To account for the high error-rate and relatively low germline coverage of only ~15x, a three-step phasing approach was developed. **(A)** In step one, single-nucleotide variants and short insertions and deletions were phased using Whatshap[1], yielding an N50 phased block length of 2.29Mbp for chromosome 3. Each dot is a phased heterozygous variant colored by haplotype with the tumor variant allele frequency on the y-axis. **(B)** Statistical phasing using ShapeIt[2] with the long-read phased blocks as input increased the N50 phased block length to 4.68Mbp. **(C)** The unequal haplotype dosage of chromosome 3 in the primary tumor (haploid p-arm, triploid q-arm) was used to correct remaining switch errors on chromosome 3.

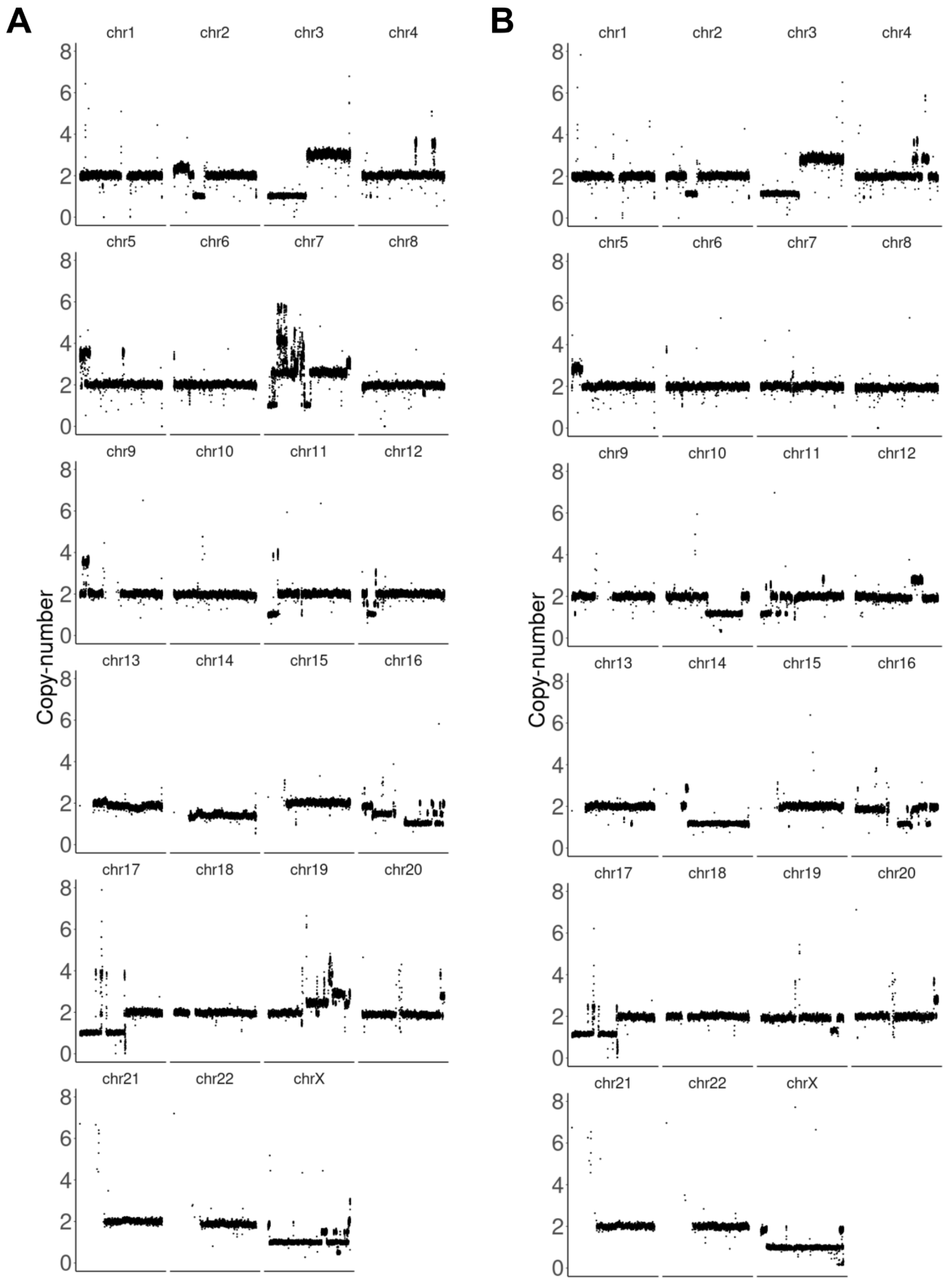


**Figure S3. Copy-number profiles.** The normalized copy-number (y-axis) of each chromosome (x-axis) using a 10kbp window length of uniquely mappable positions and GC-fragment corrected read counts from delly's cnv mode[3]. Panel **(A)** shows the primary tumor and panel **(B)** the relapse sample.

**A**

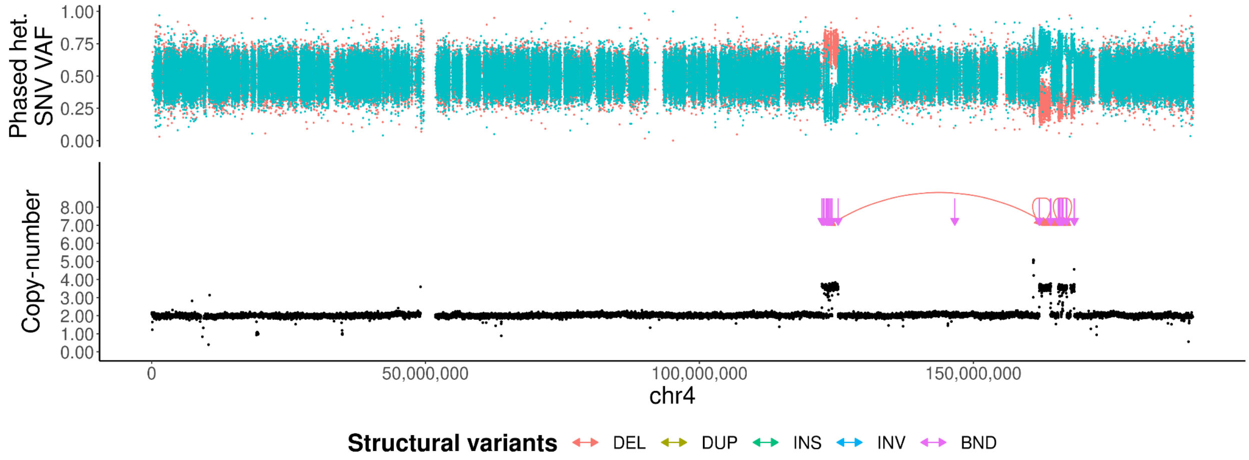
Phased het. SNV VAF
1.00
0.75
0.50
0.25
0.00
Copy-number
8.00
7.00
6.00
5.00
4.00
3.00
2.00
1.00
0.00
0
50,000,000
100,000,000
150,000,000
chr4
Structural variants
DEL
DUP
INS
INV
BND

**B**

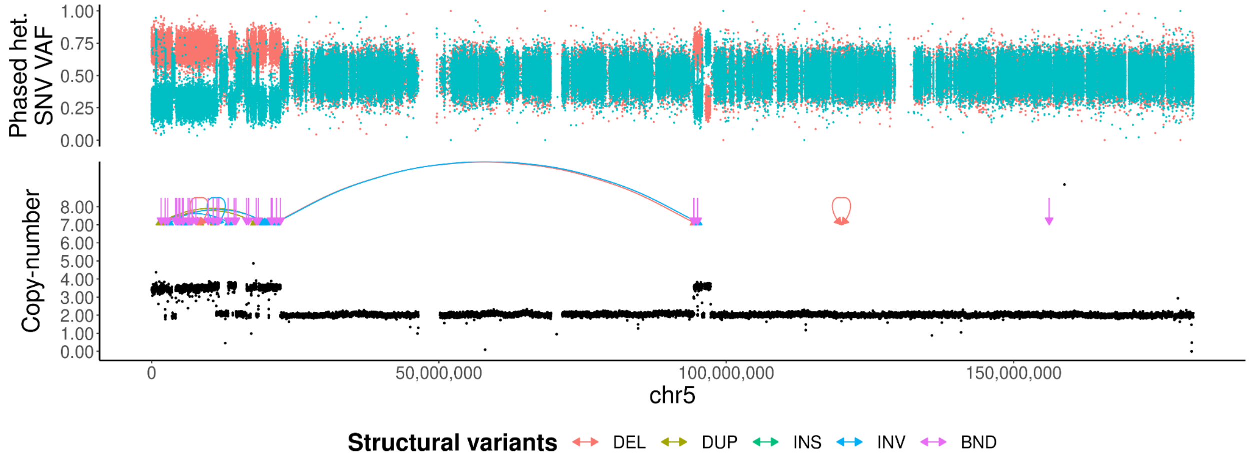
Phased het. SNV VAF
1.00
0.75
0.50
0.25
0.00
Copy-number
8.00
7.00
6.00
5.00
4.00
3.00
2.00
1.00
0.00
0
50,000,000
100,000,000
150,000,000
chr5
Structural variants
DEL
DUP
INS
INV
BND

**C**

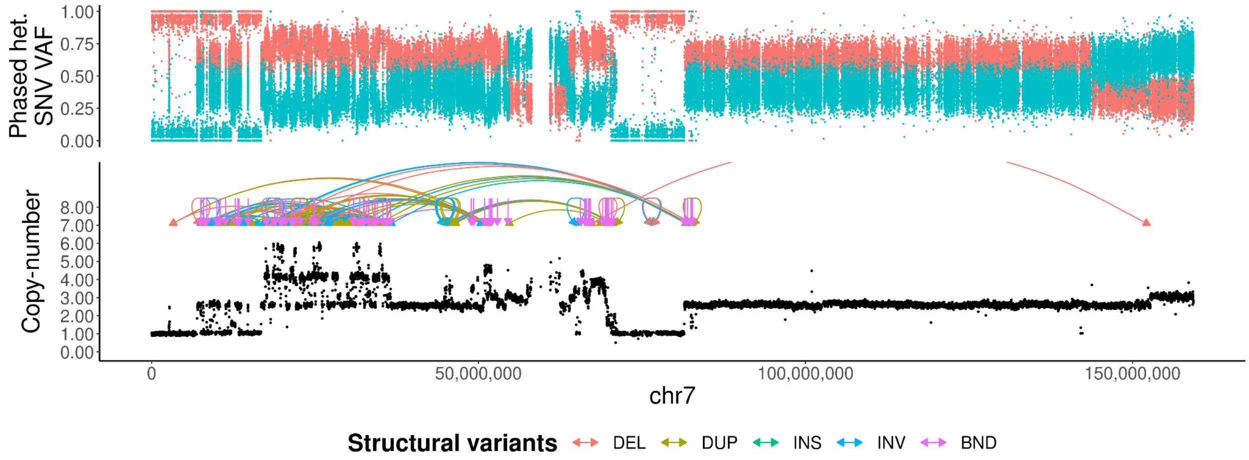
Phased het. SNV VAF
1.00
0.75
0.50
0.25
0.00
Copy-number
8.00
7.00
6.00
5.00
4.00
3.00
2.00
1.00
0.00
0
50,000,000
100,000,000
150,000,000
chr7
Structural variants
DEL
DUP
INS
INV
BND

**D**

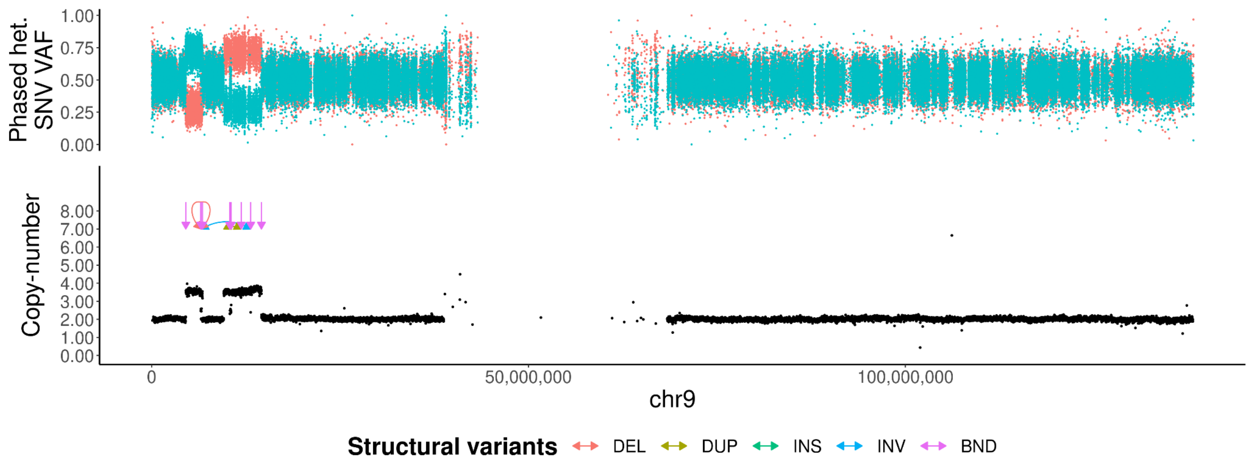
Phased het. SNV VAF
Copy-number
0
50,000,000
100,000,000
chr9
Structural variants
DEL
DUP
INS
INV
BND

**E**

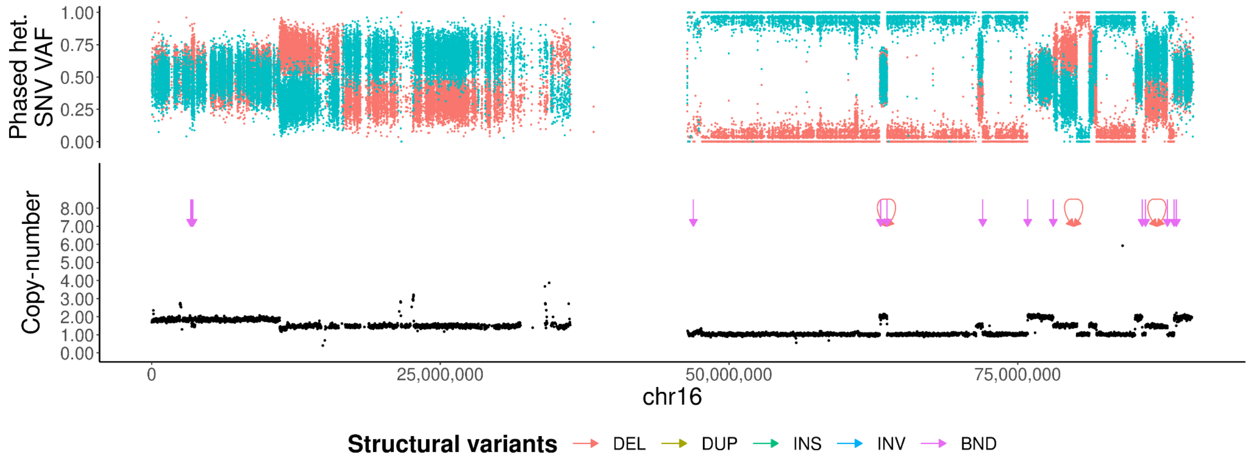
Phased het. SNV VAF
Copy-number
0
25,000,000
50,000,000
75,000,000
chr16
Structural variants
DEL
DUP
INS
INV
BND

**F**

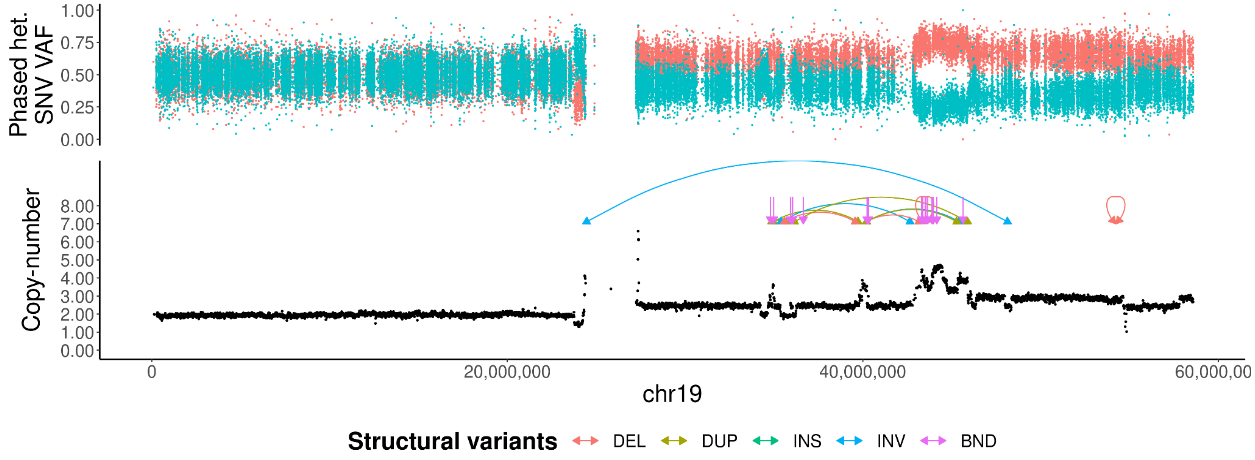
Phased het. SNV VAF
Copy-number
0
20,000,000
40,000,000
60,000,00
chr19
Structural variants
DEL
DUP
INS
INV
BND

**G**

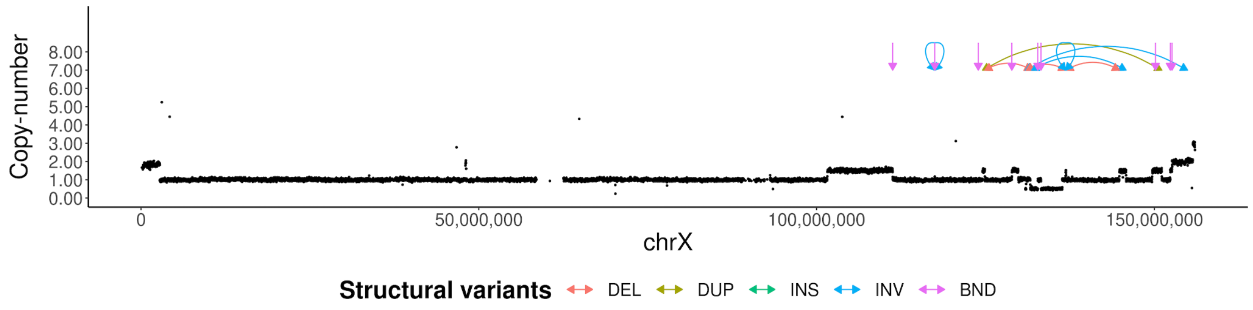


**Figure S4. Copy-number plots of chromothripsis chromosomes.** The upper panel shows the single-nucleotide variant (SNV) allele frequency (VAF) for phased heterozygous germline variants colored by haplotype. The lower panel shows the normalized copy-number (y-axis) of each chromosome (x-axis) using a 10kbp window length. Overlaid are somatic structural variant (SV) calls colored by SV type. Panel **(A)** shows chromosome 4, **(B)** shows chromosome 5, **(C)** shows chromosome 7, **(D)** shows chromosome 9, **(E)** shows chromosome 16, **(F)** shows chromosome 19 and panel **(G)** shows the haploid chromosome X (male sample).

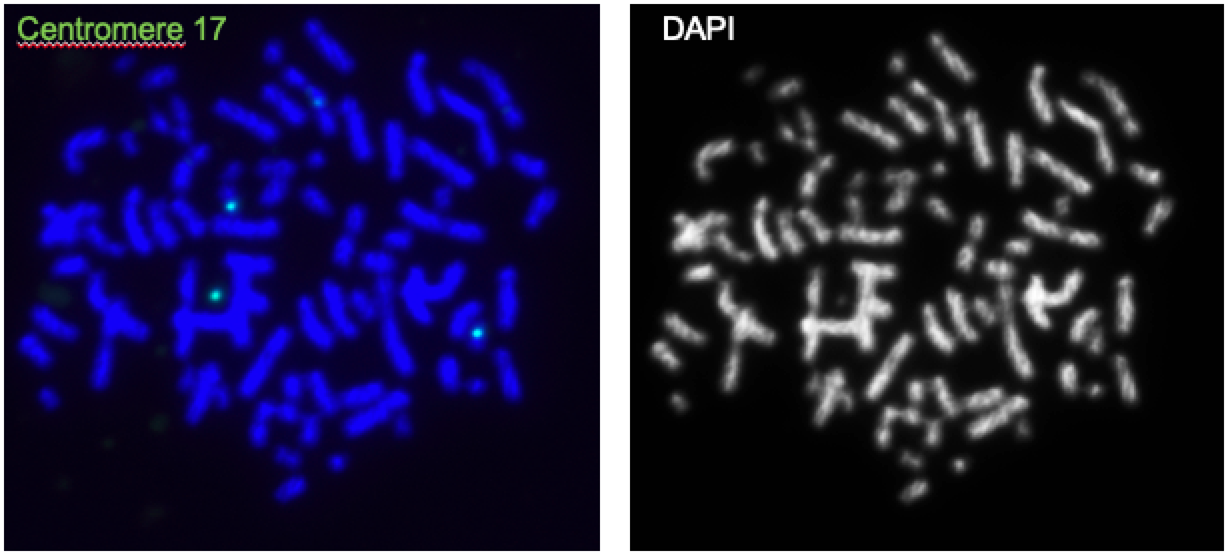


**Figure S5. FISH analysis on metaphase spreads from patient-derived xenografted cells.** Signals for the centromere 17 probe (left) and DAPI (right) suggest an absence of double-minute chromosomes. From the three centromere signals detected here, two are on chromosomes and one is on a chromosome fragment (lower left).

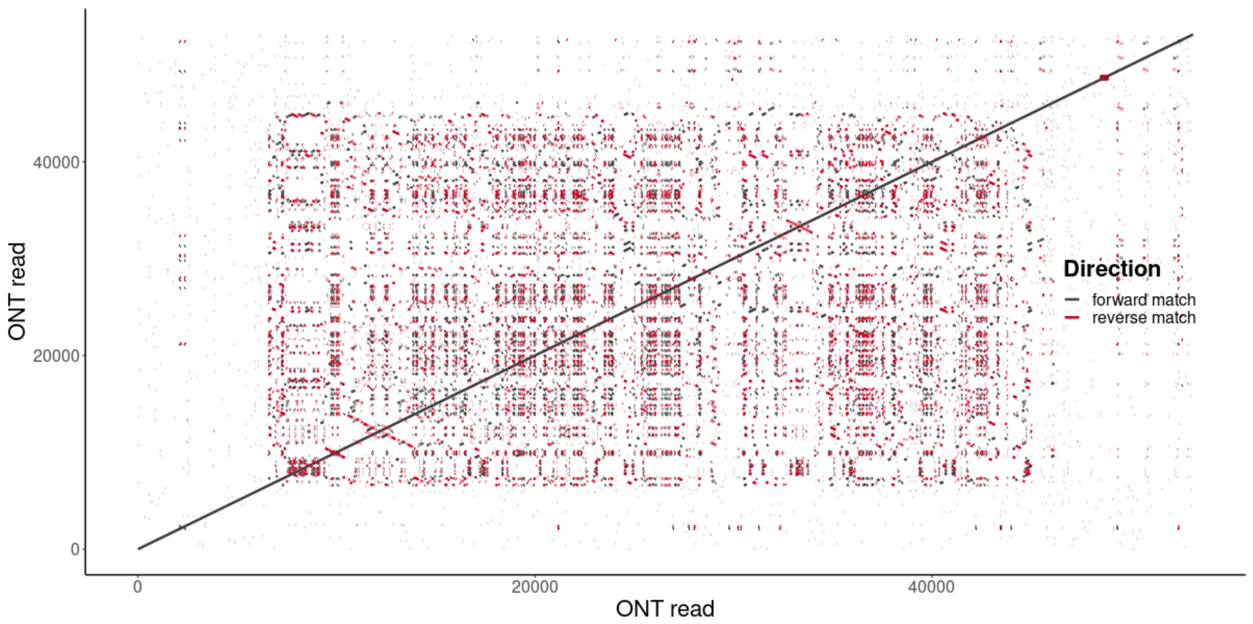


**Figure S6. Self-alignment of the templated insertion thread.** Forward and reverse matches of a single ONT read aligned against itself for the 2nd instance of a templated insertion thread in the primary tumor, as in Figure 2A for the 1st instance.

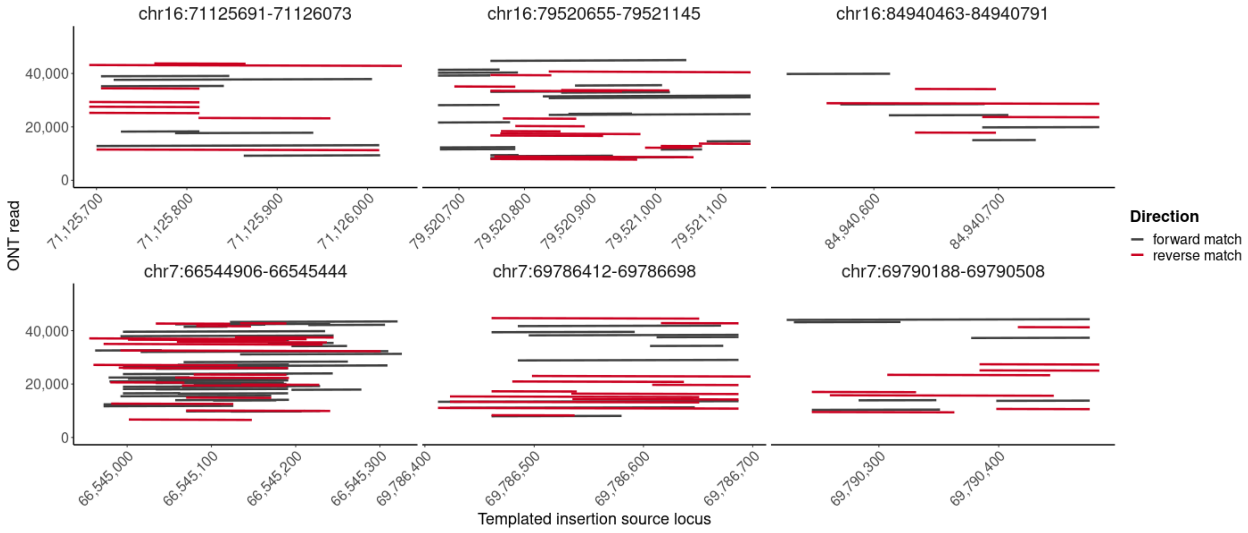


**Figure S7. Templated insertion thread alignments.** Similar to Figure 2B, forward and reverse matches for a subset of the templated insertion source sequences of the 2nd instance of a templated insertion thread.

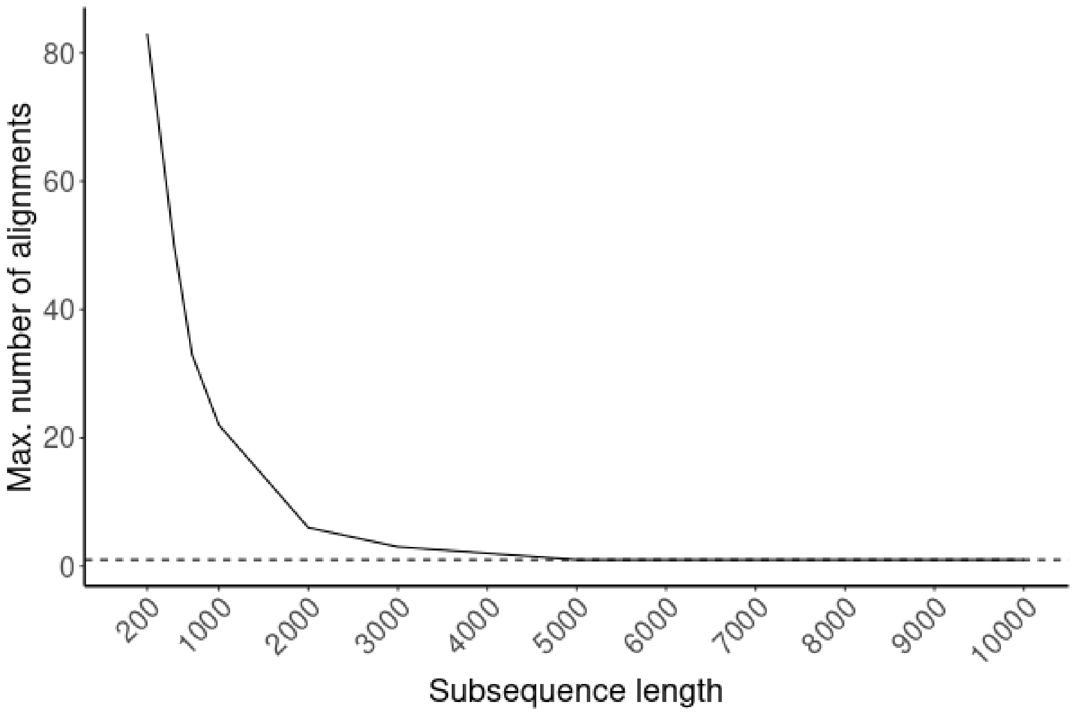


**Figure S8. Subsampling analysis of ONT reads spanning templated insertion threads.** To identify possible higher-order repeating structures, we randomly sampled sub-sequences of different length (x-axis). Each sub-sequence was iteratively aligned to the original ONT read, masking all previous alignments until no further confident alignments could be identified. The dashed line corresponds to the original sampling location without any further alignments (y=1).

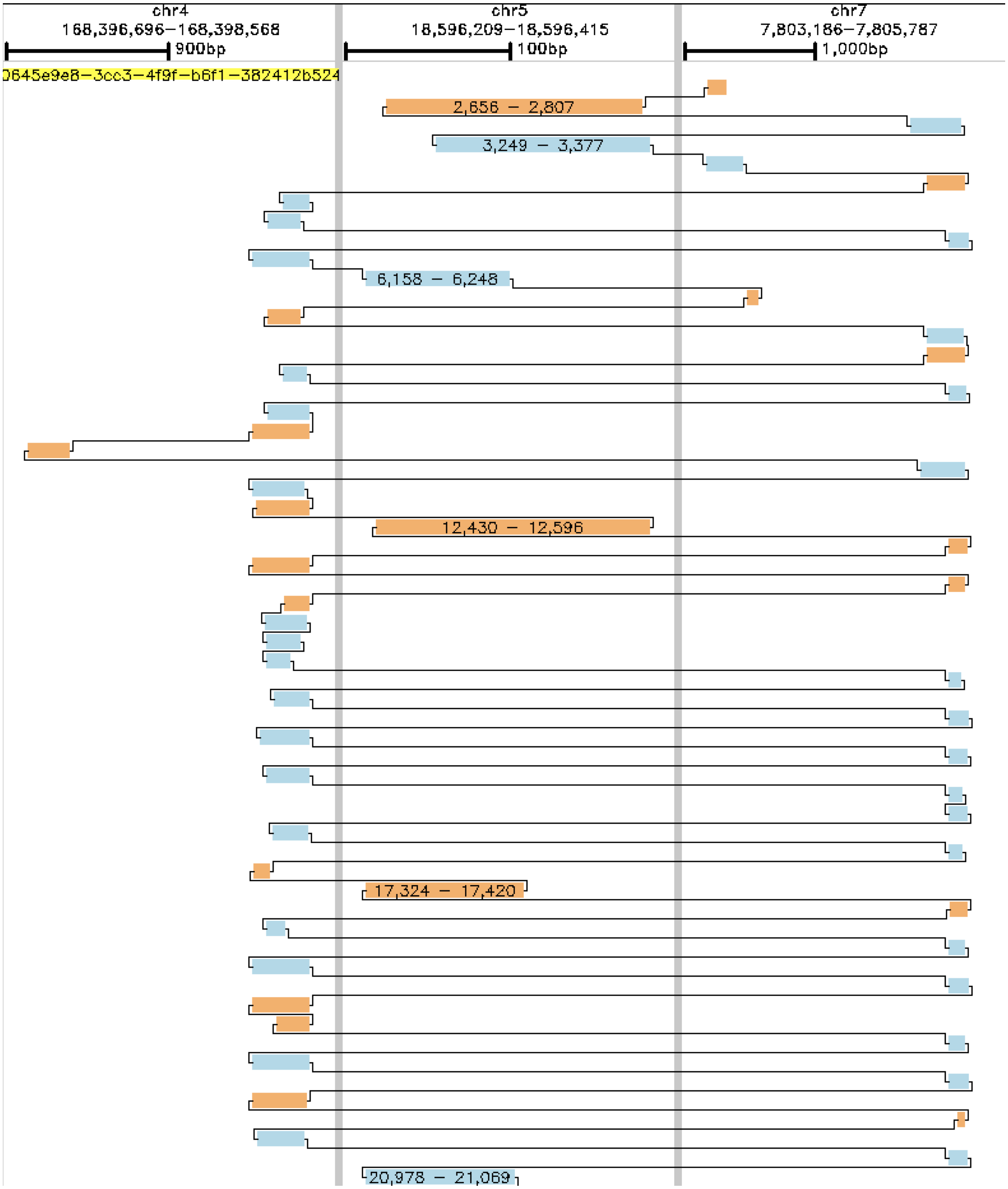


**Figure S9. Genomic matches of a single ONT read with a templated insertion thread.** Forward matches are in blue, reverse matches in orange and the black line traces the matches along the read, concatenating in a zig-zag fashion the templated insertion source sequences. Each vertical panel is a separate genomic alignment region specified in the header. Only a subset of the matches in the first 22kbp of the read are shown for illustration reasons, the full alignment of this read is available as a separate file: 0645e9e8-3cc3-4f9f-b6f1-382412b524b6.png .

**A) Cluster 1**

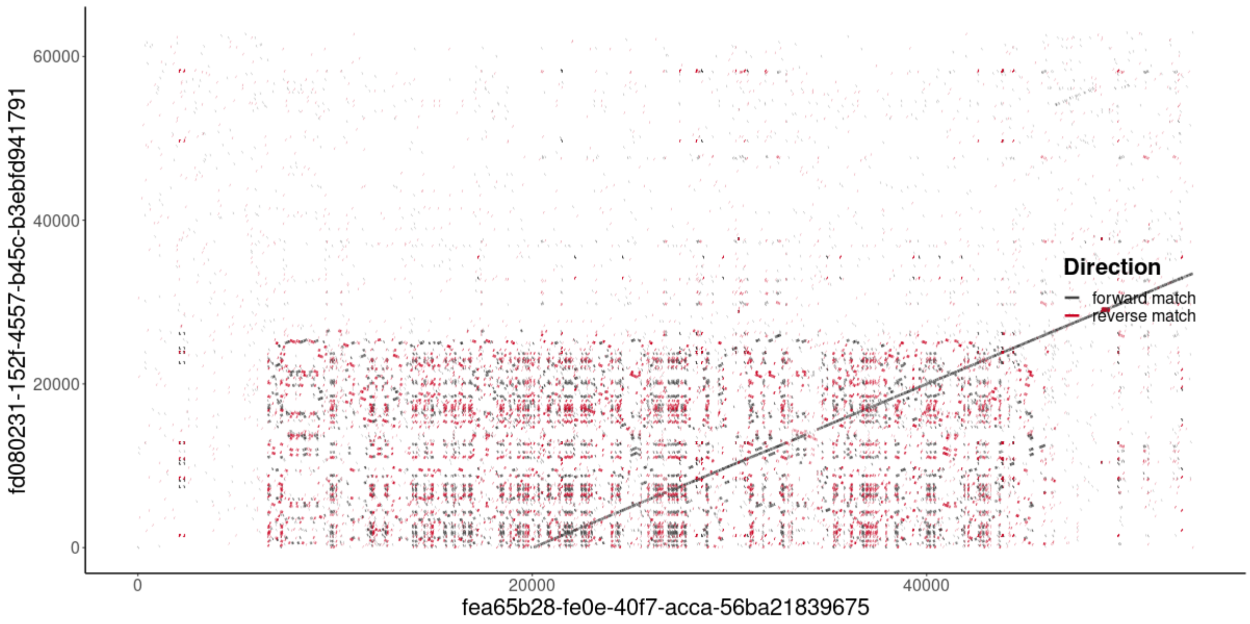


**B) Cluster 2**

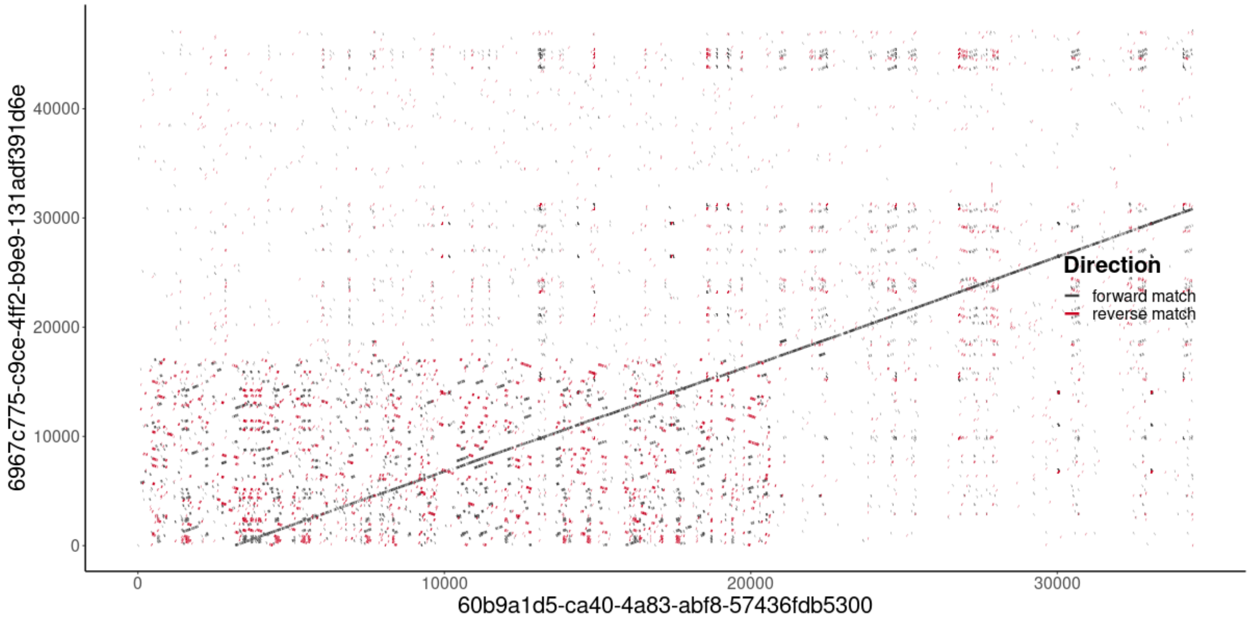


**C) Cluster 3**

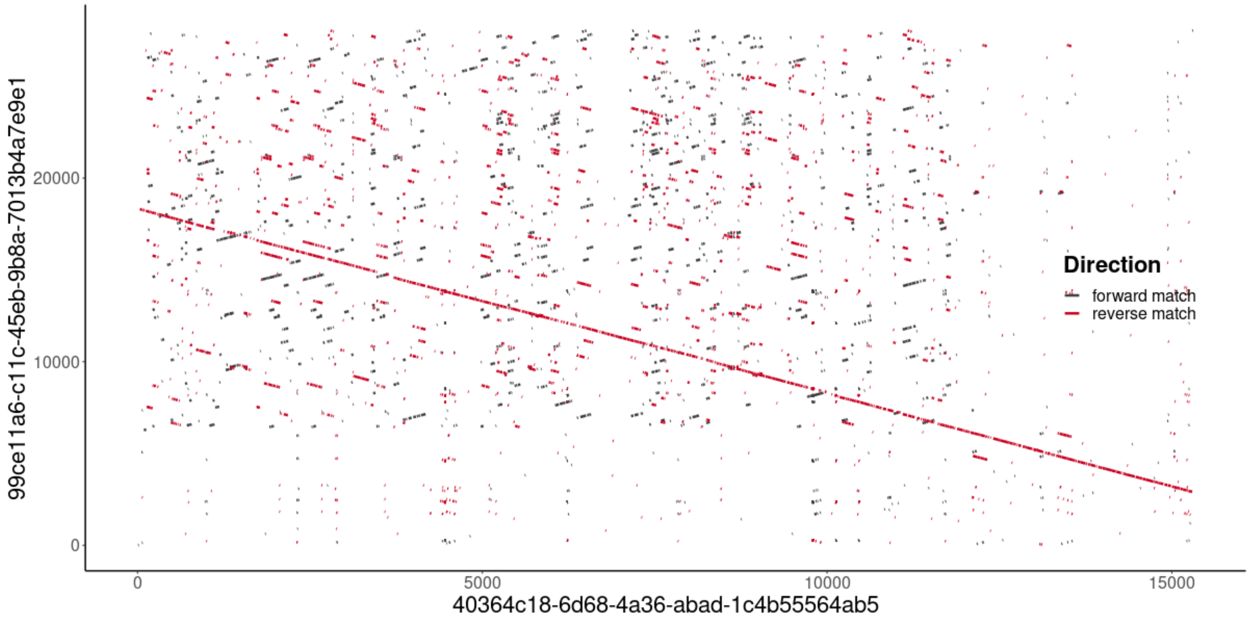


**D) Genome alignments to templated insertion source segments**

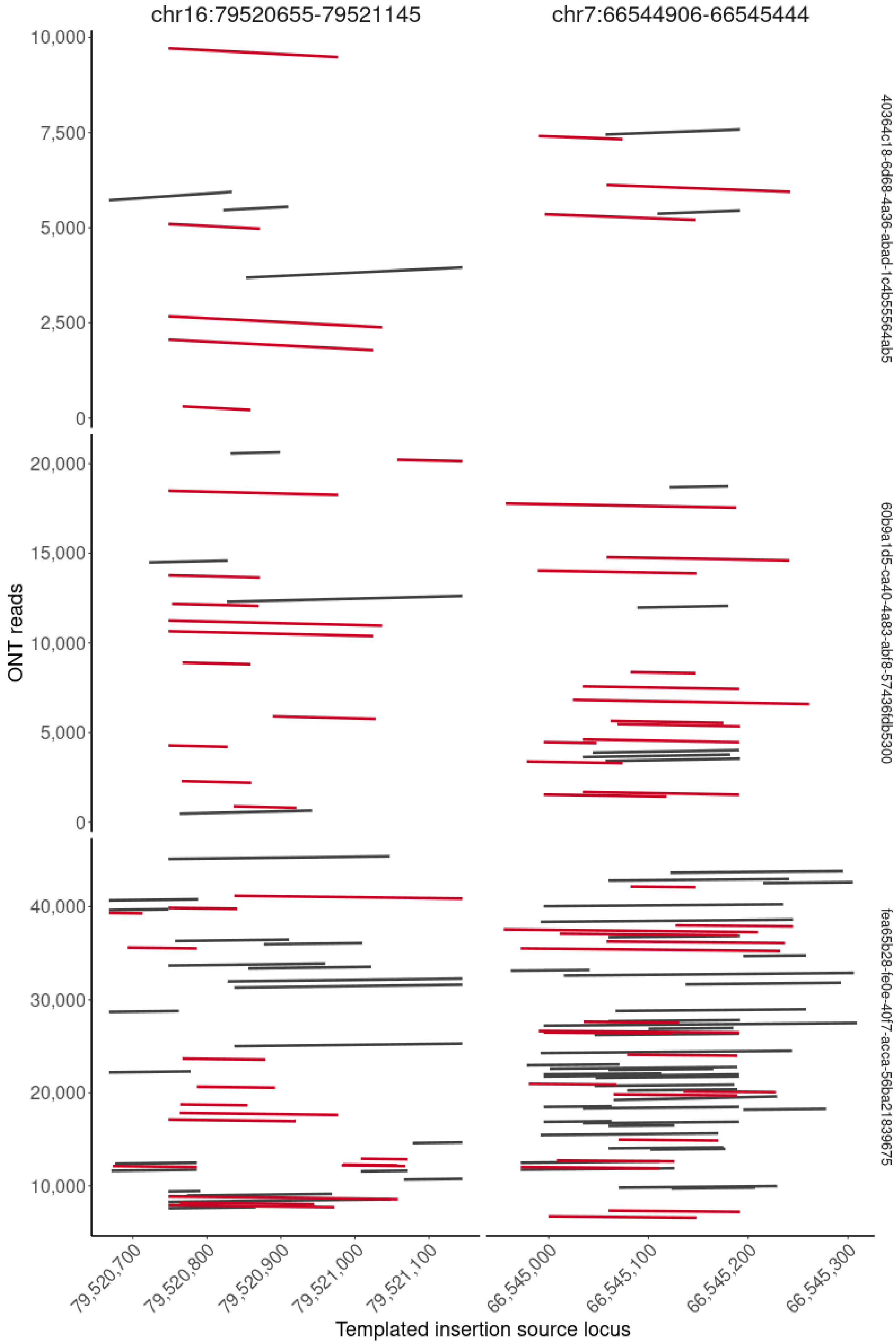
chr16:79520655-79521145
chr7:66544906-66545444
ONT reads
Templated insertion source locus
40364c18-6d68-4a36-abad-1c4b55564ab5
60b9a1d5-ca40-4a83-abf8-57436fdb5300
fea65b28-fe0e-40f7-acca-56ba21839675
10,000
7,500
5,000
2,500
0
20,000
15,000
10,000
5,000
0
40,000
30,000
20,000
10,000
79,520,700
79,520,800
79,520,900
79,521,000
79,521,100
66,545,000
66,545,100
66,545,200
66,545,300

**Figure S10. Tumor heterogeneity of templated insertion threads.** Templated insertion threads show signals of tumor heterogeneity. **(A-C)** Pairwise alignments of two ONT reads supporting the same templated insertion thread architecture with **(D)** showing large differences among these three clusters of reads by means of how often the same templated insertion source segments are concatenated and in what order and orientation.

**A**

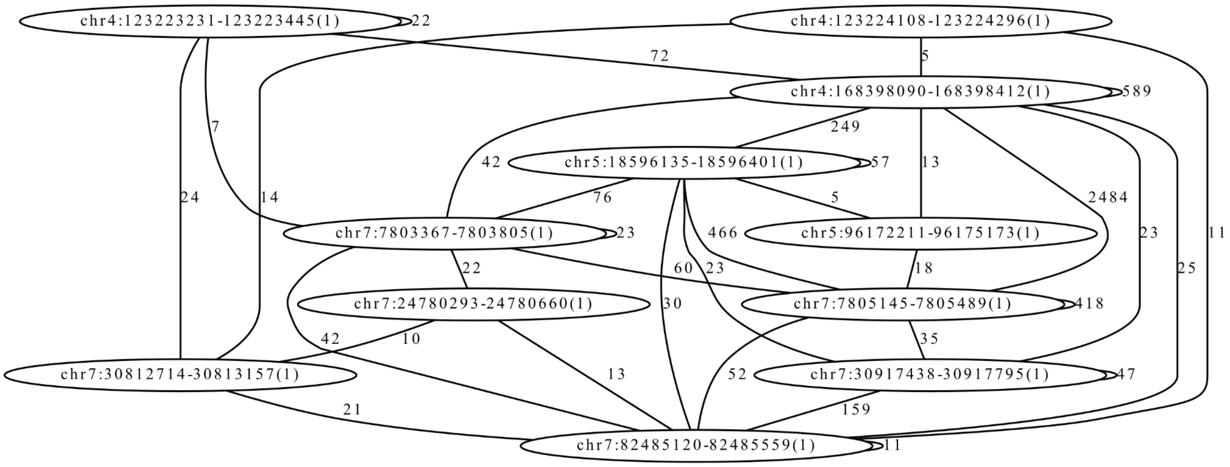


**B**

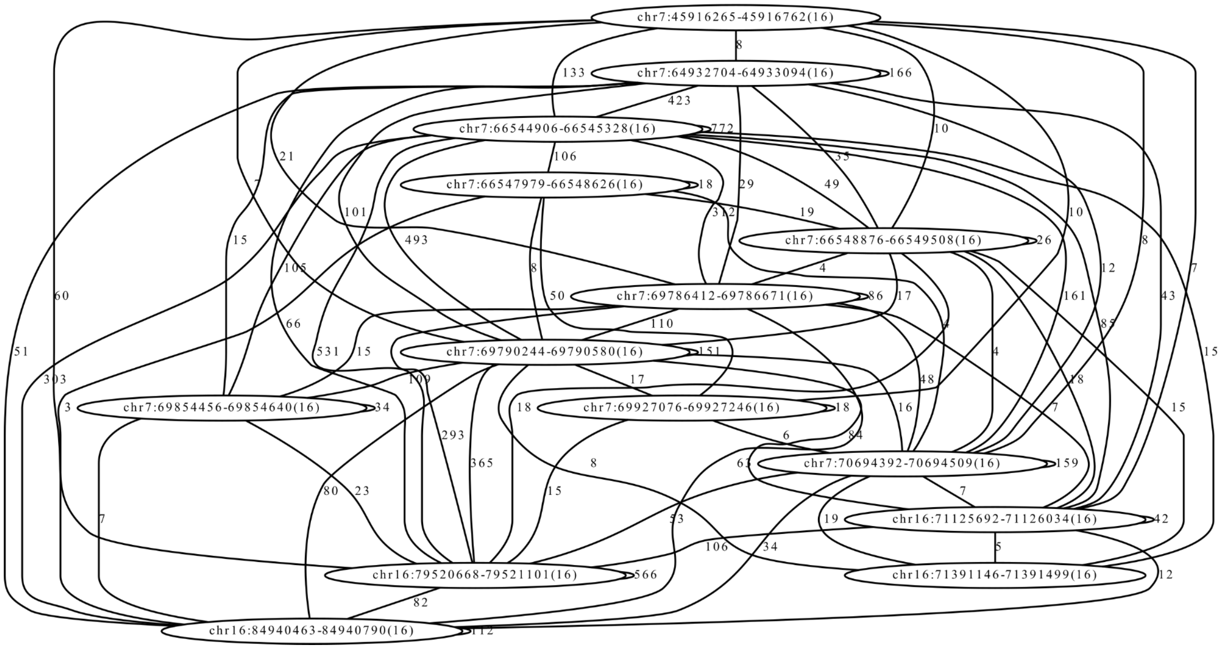


**Figure S11. Templated insertion rearrangement graphs.** Nodes of the graph represent templated insertion source sequences. Edges of the graph represent split-read and paired-end connections between different templated insertions, including self edges. The edge weight specifies the support for the rearrangement junction. The 1st instance in panel **(A)** corresponds to Figure 2AB and the 2nd instance in panel **(B)** corresponds to Figure S6 and S7.

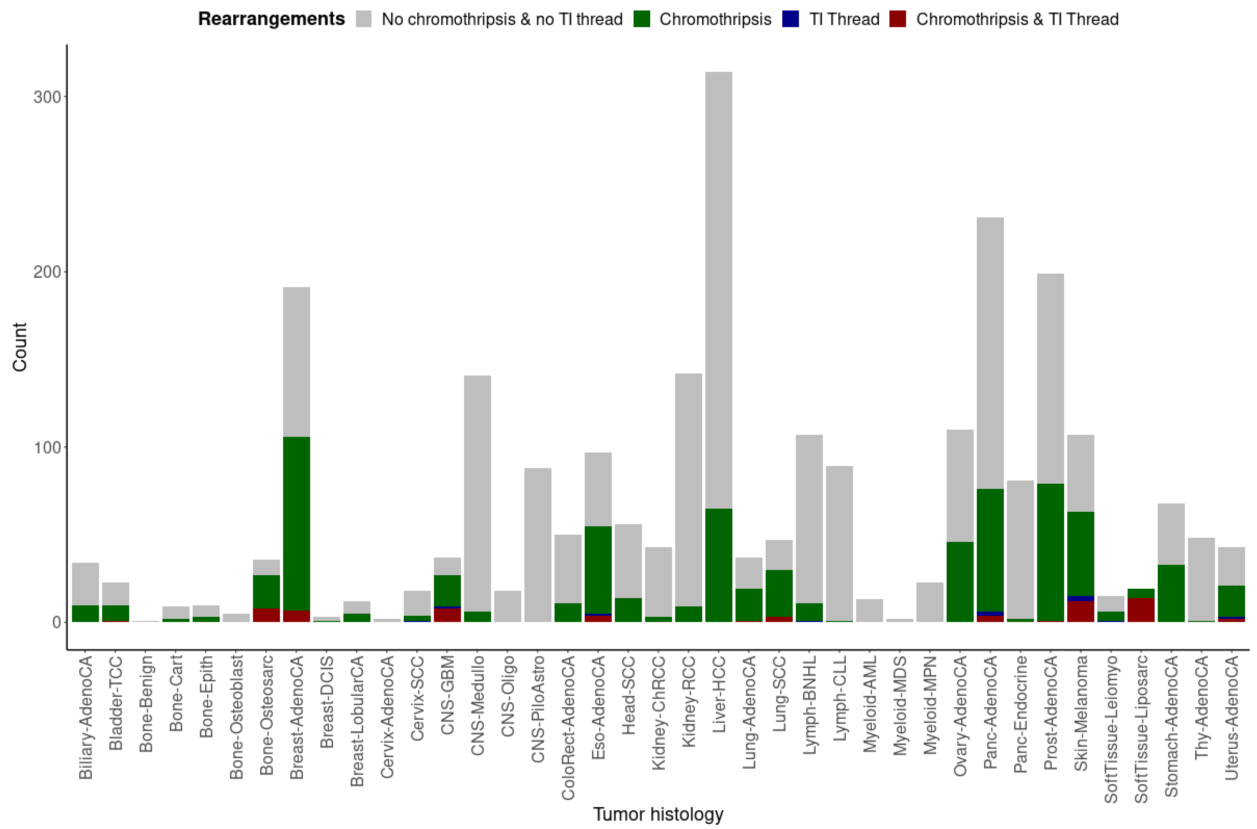


**Figure S12. Templated insertion threads across 2,569 cancer genomes.** A stacked histogram of the distribution of templated insertion threads (TI Threads) and chromothripsis across all PCAWG samples stratified by tumor histology.

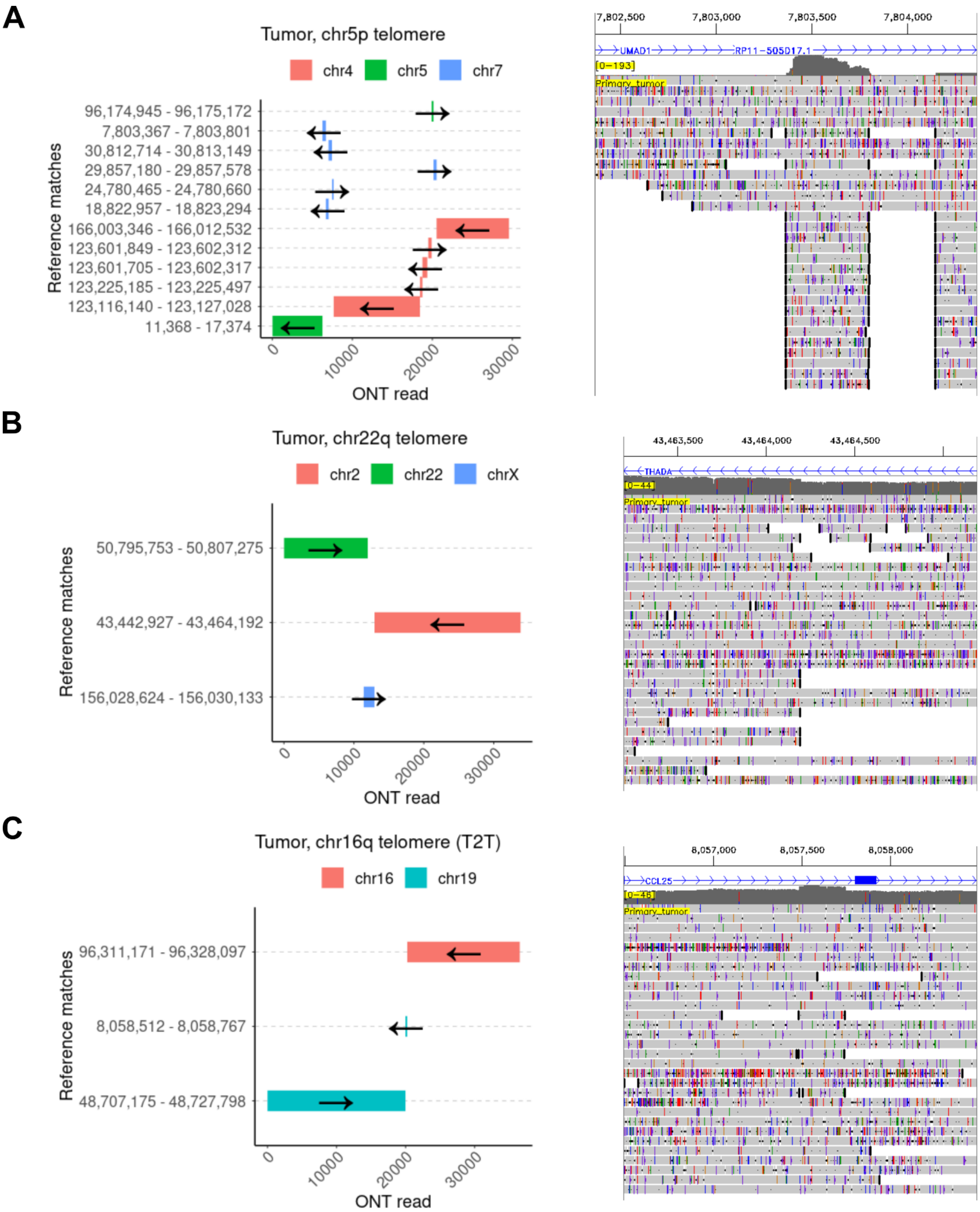
A
Tumor, chr5p telomere
chr4
chr5
chr7
96,174,945 - 96,175,172
7,803,367 - 7,803,801
30,812,714 - 30,813,149
29,857,180 - 29,857,578
24,780,465 - 24,780,660
18,822,957 - 18,823,294
166,003,346 - 166,012,532
123,601,849 - 123,602,312
123,601,705 - 123,602,317
123,225,185 - 123,225,497
123,116,140 - 123,127,028
11,368 - 17,374
Reference matches
0
10000
20000
30000
ONT read
7,802,500
7,803,000
7,803,500
7,804,000
UMAD1
RP11-505D17.1
[0-193]
Primary_tumor
B
Tumor, chr22q telomere
chr2
chr22
chrX
50,795,753 - 50,807,275
43,442,927 - 43,464,192
156,028,624 - 156,030,133
Reference matches
0
10000
20000
30000
ONT read
43,463,500
43,464,000
43,464,500
THADA
[0-44]
Primary_tumor
C
Tumor, chr16q telomere (T2T)
chr16
chr19
96,311,171 - 96,328,097
8,058,512 - 8,058,767
48,707,175 - 48,727,798
Reference matches
0
10000
20000
30000
ONT read
8,057,000
8,057,500
8,058,000
CCL25
[0-46]
Primary_tumor

**D**

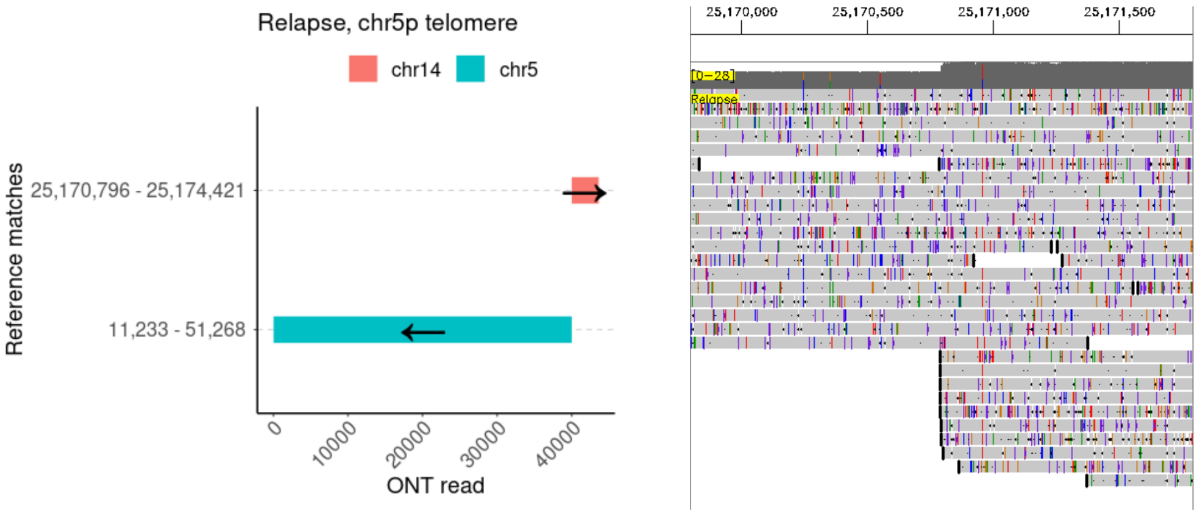


**E**

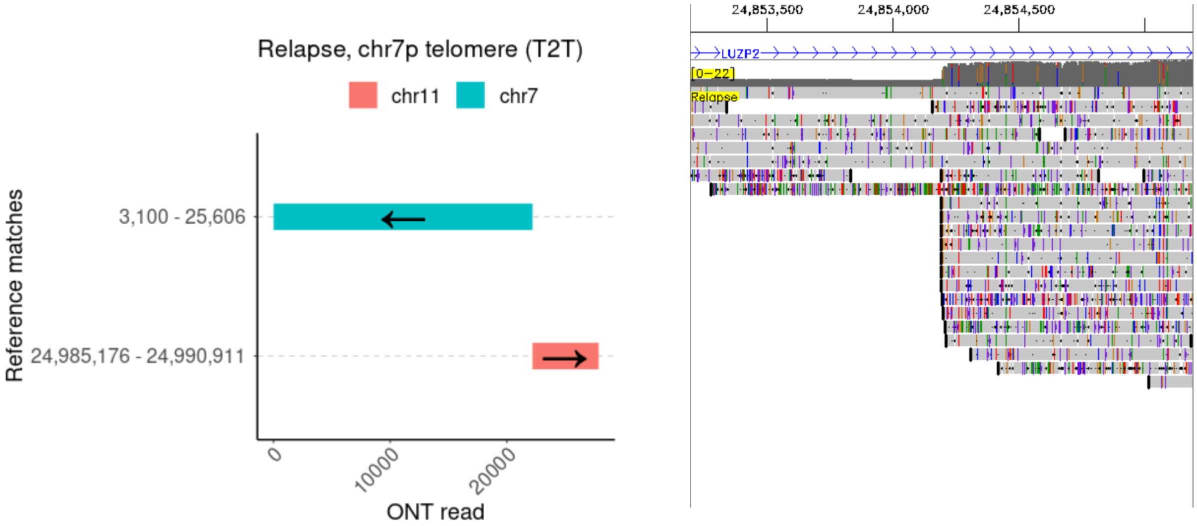


**Figure S13. Telomere sequences associated with rearranged genomic regions.** Panels (**A-E**) show five telomere sequences associated with SVs, three in the primary tumor (**A-C**) and two in relapse (**D-E**), using a representative ONT read (x-axis) and the genomic mapping locations (y-axis) colored by chromosome (left panels). Coordinates are in GRCh38, except for rearrangements mapped to the telomere to telomere assembly (T2T) to resolve alignment ambiguities (panel C and E). The panels on the right show a selected non-telomeric SV breakpoint spanned by the respective read from the left panel. Some of these breakpoints are within an intron of a gene (panel A, B, C, and E) or show alignment signals characteristic for templated insertions (panel A and C).

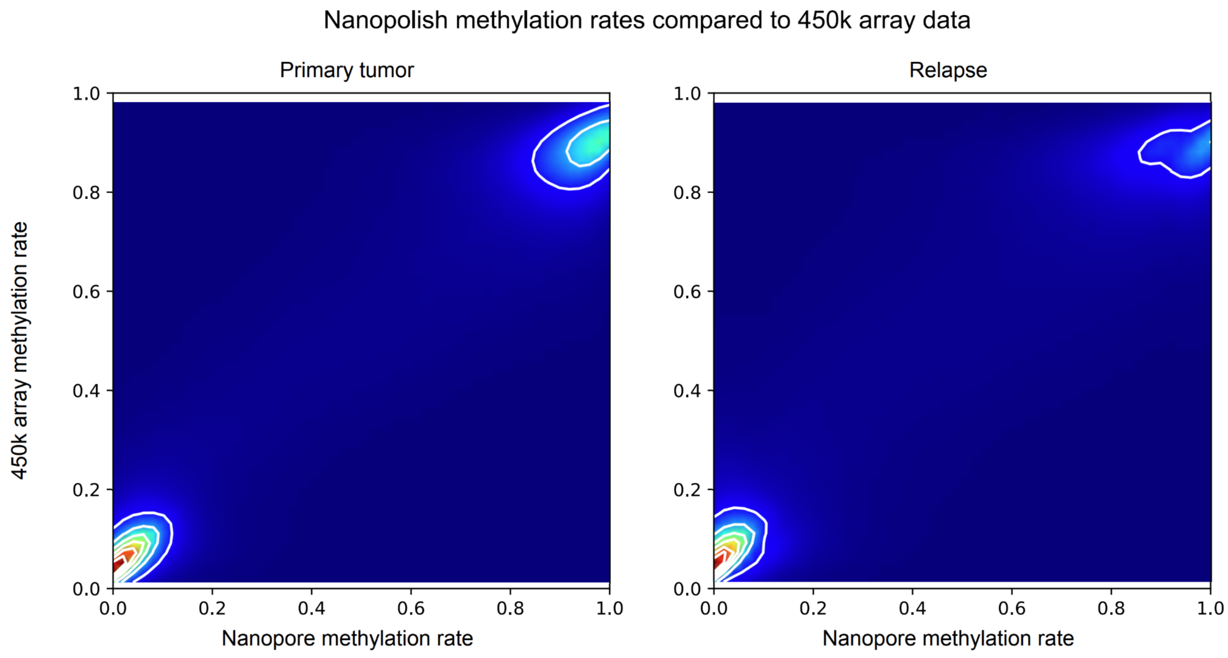


**Supplementary Figure S14.** Validation of nanopolish methylation calls using Illumina 450k array data. Plotting each array probe's methylation rate against the mean methylation rate in the matching region from nanopolish methylation calls (log-likelihood ratio threshold 2.0). Pearson correlation r=0.9540 for primary tumor and r=0.9218 for the relapse sample.

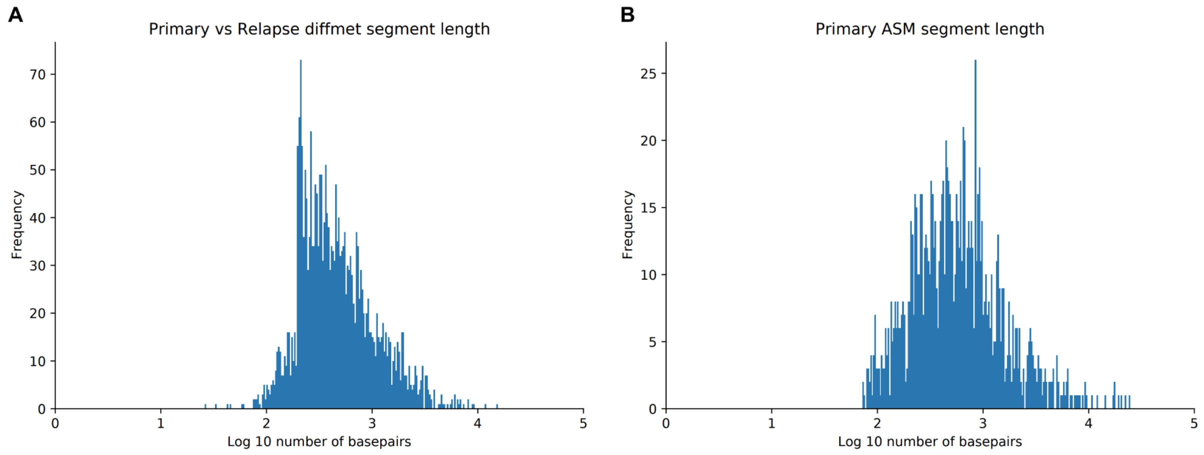


**Supplementary Figure S15.** Length of differentially methylated segments (A) in the primary tumor vs relapse sample comparison and (B) allele specific methylation analysis in the primary tumor sample.

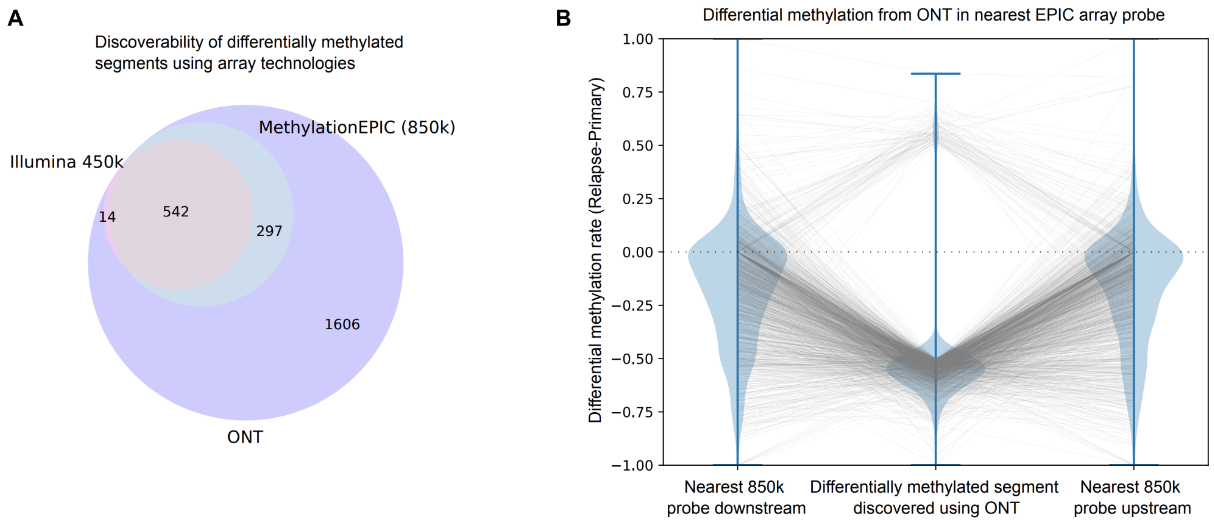


**Supplementary Figure S16.** Differentially methylated regions between primary tumor and relapse sample discovered from ONT (abs methylation rate difference > 0.5) have been tested for discoverability using Illumina 450k array and MethylationEPIC (850k) array technologies. A) Counting the number of segments for which an array probe at least partially overlapped a segment found as differentially methylated in ONT. B) For the 1620 segments not covered by a probe in the MethylationEPIC array, we investigate the methylation as measured from ONT at the location of the nearest array probes upstream and downstream from the differentially methylated segment. We find that only for 386 segments one of the two probes shows matching differential methylation.

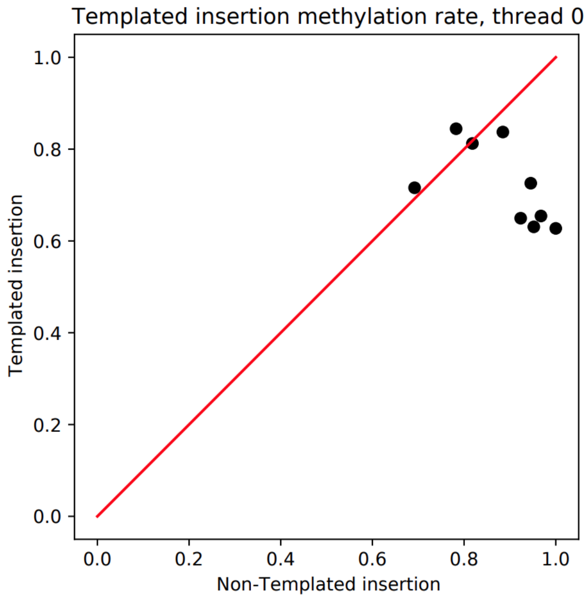


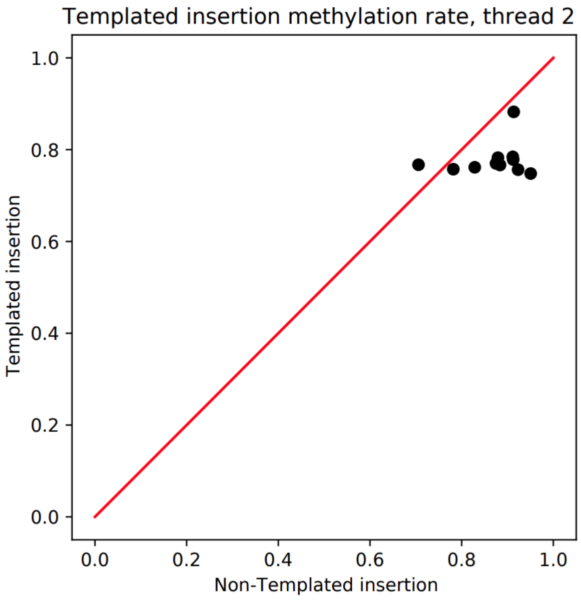


**Supplementary Figure S17.** A number of high coverage templated insertions from two templated insertion threads. Methylation rates from templated insertions are plotted against methylation rates from the unmodified haplotype at the insertions location of origin. Only a minor reduction in methylation rate (by about 0.16 in thread 1 and about 0.09 in thread 2) can be observed in templated insertions.

A)

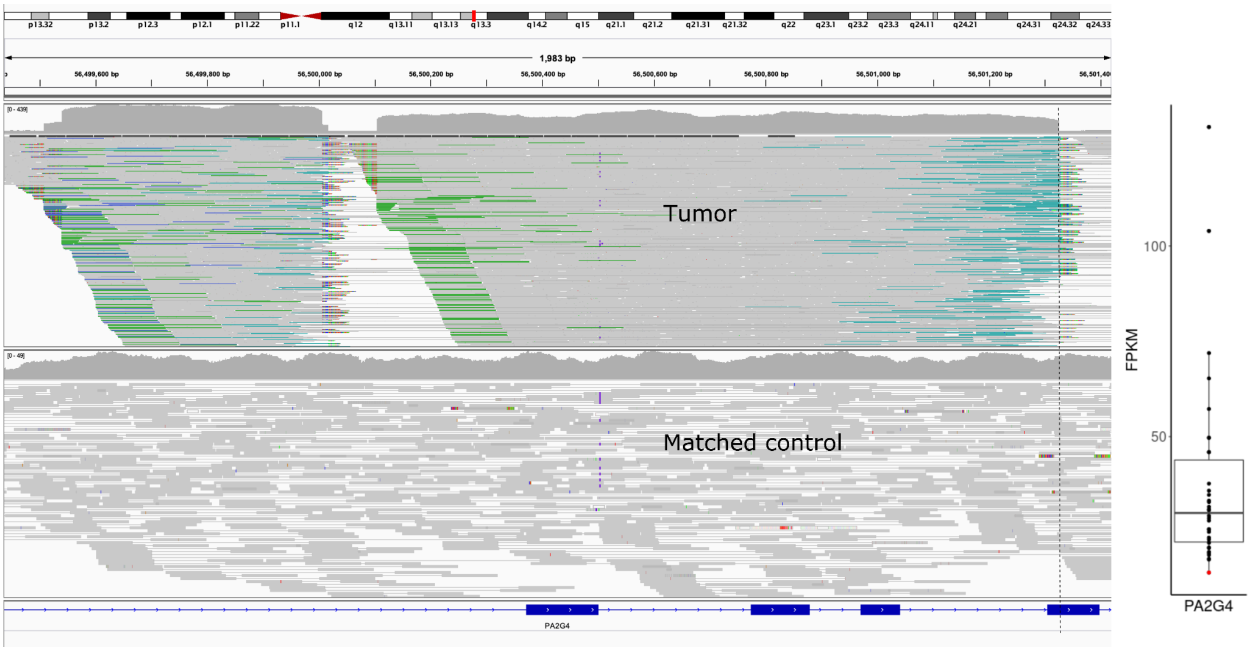


B)

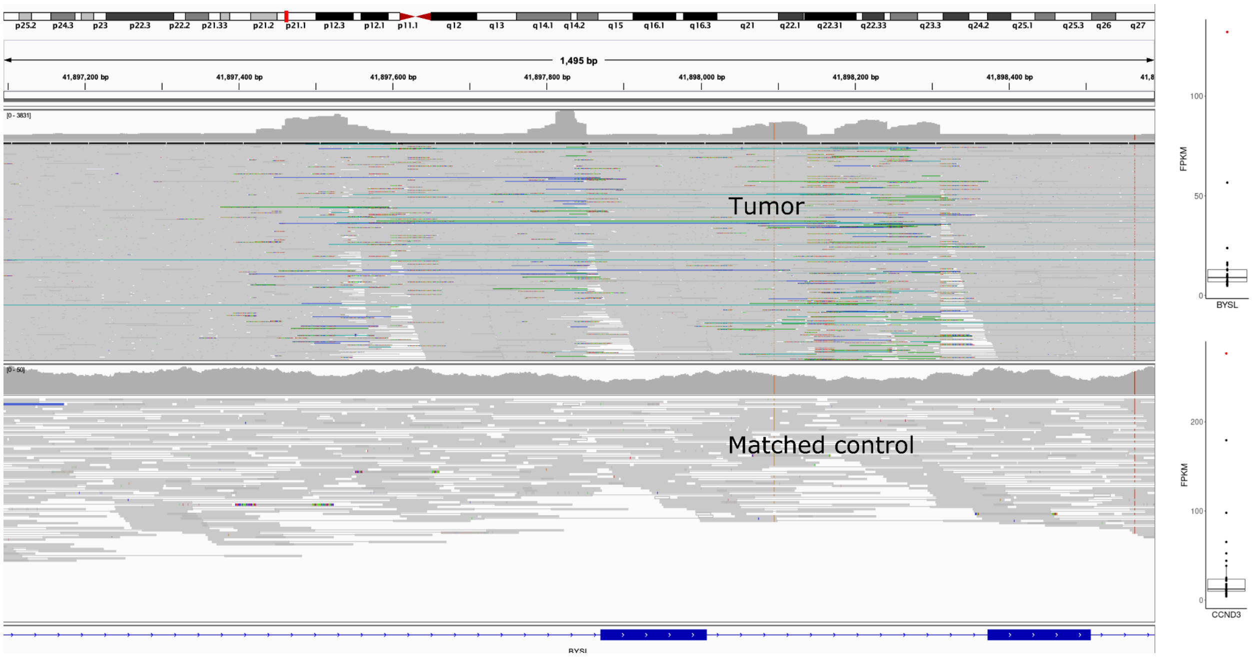


**Figure S18. Gene expression effects of templated insertion threads. (A)** Example of a templated insertion thread identified by *rayas* that intersects a coding exon of PA2G4. The boxplot on the right shows the FPKM distribution of all sarcoma samples in PCAWG with the tumor shown on the left having the lowest expression among all samples, likely due to the exon intersecting templated insertion thread. **(B)** Overexpression of BYSL and CCND3 relative to other sarcoma samples (right boxplots) and the alignment view of a templated insertion thread overlapping multiple introns of BYSL (left alignment view).

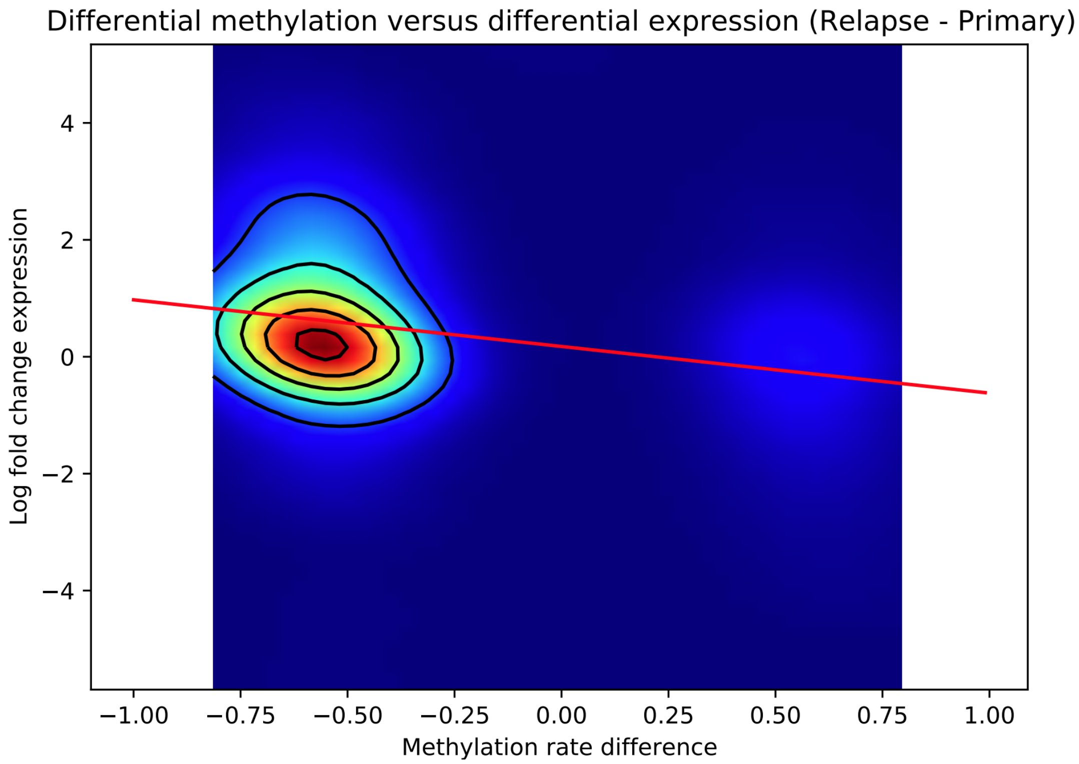


**Supplementary Figure S19.** Differential promoter methylation between primary tumor and relapse shows significant inverse correlation with gene expression log-fold change (Spearman-R -0.31, p-value: $1.8 \times 10^{-2}$), showing promoter methylation driven transcription.

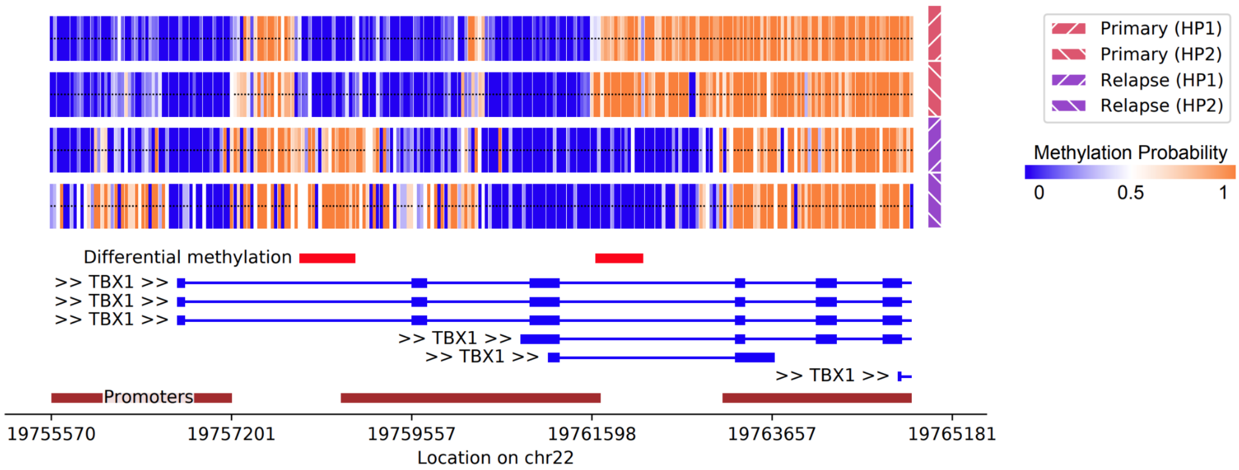


**Supplementary Figure S20.** Two promoter linked DMRs in TBX1 which is also highly expressed in the relapse sample. The DMR demethylated in primary tumor precedes the short form transcript TBX1-206, whereas the DMR highly methylated in primary tumor directly succeeds the transcription start site of said transcript.

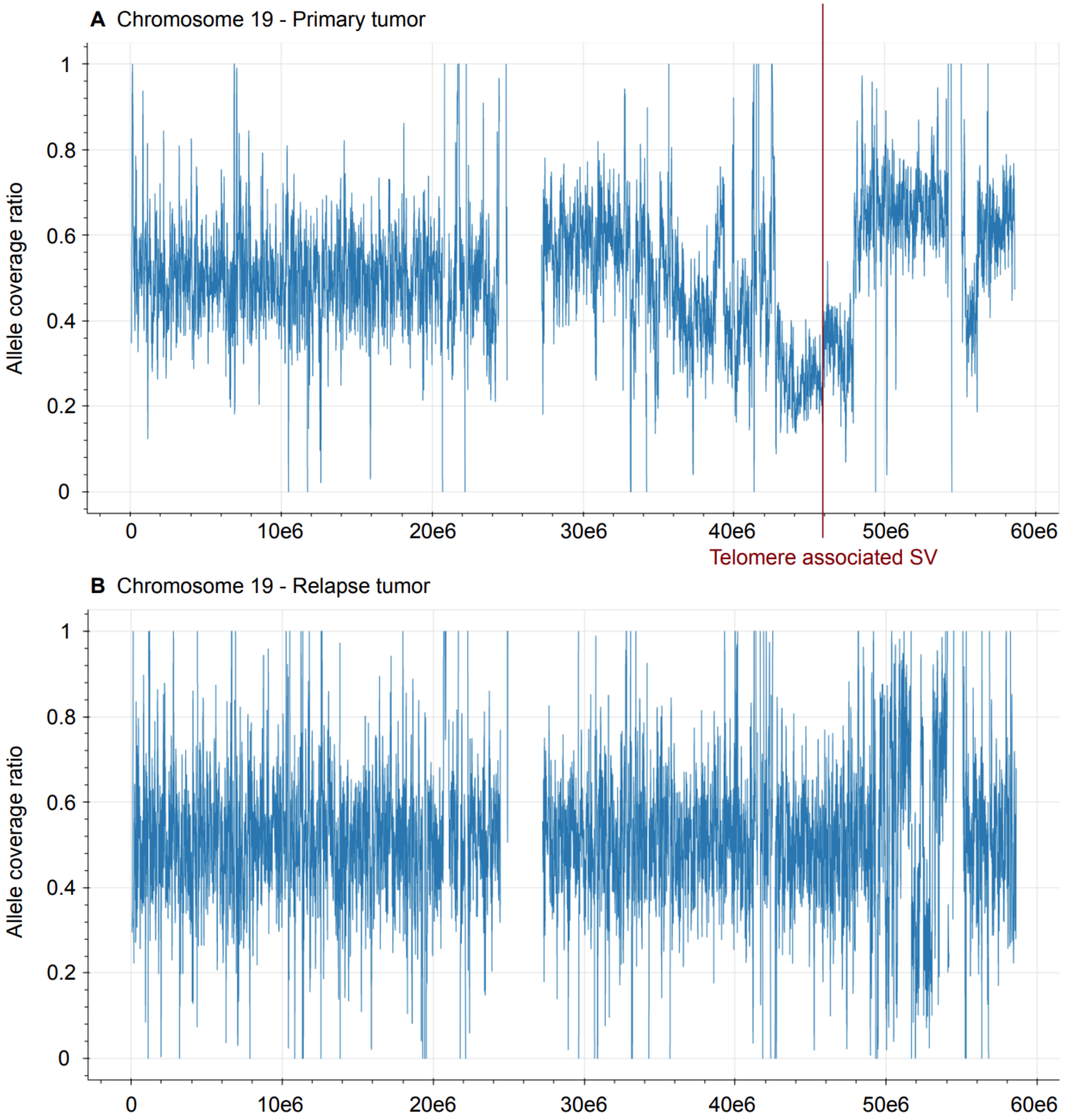


**Supplementary Figure S21.** Allele coverage ratio from WhatsHap phased ONT reads of chromosome 19 in primary (**A**) and relapse (**B**) tumor. The telomere associated SV observed in primary tumor coincides with a haplotype specific amplification of chromosome 19q.

### Supplementary Tables

**Table S1**. Long-read sequencing statistics

| | **Germline** | **Primary tumor** | **Relapse** |
|---|---|---|---|
| **Coverage** | 15x | 30x | 15x |
| **Median read length** | 4,480bp | 4,993bp | 5,678bp |
| **Est. error rate** | 8.5% | 6.8% | 6.9% |

**Table S2**. Short-read sequencing statistics

| | **Germline** | **Primary tumor** | **Relapse** |
|---|---|---|---|
| **Coverage** | 48x | 45x | 47x |
| **Seq. mode** | 2 * 150bp | 2 * 150bp | 2 * 150bp |
| **Insert size** | 373bp | 387bp | 406bp |

**Table S3**. FISH analysis using probes for RP11-651L9 and centromere 17 (primary tumor).

| **Number of signals** | **651L9** | **Centr.17** |
|---|---|---|
| 0 | 2 | 0 |
| 1* | 23 | 4 |
| 2** | 36 | 20 |
| 3 | 20 | 30 |
| 4 | 16 | 28 |
| 5 | 3 | 15 |
| 6 | | 2 |
| 7 | | 1 |

*Due to the tissue cutting, nuclei with only one signal can be explained by signal truncation of tumor nuclei during the sectioning process

**Normal cell nuclei are included

**Table S4**. Combined FISH analysis using probes RP11-651L9 and centromere 17.

| **Signals** | **651L9 / Centr.17** |
|---|---|
| 0/1 | 1 |
| 0/4 | 1 |
| 1/1 | 2 |
| 1/2 | 3 |
| 1/3 | 9 |
| 1/4 | 7 |
| 1/5 | 2 |
| 2/2 | 13 |
| 2/3 | 9 |
| 2/4 | 9 |
| 2/5 | 3 |
| 2/6 | 1 |
| 2/7 | 1 |
| 3/1 | 1 |
| 3/2 | 2 |
| 3/3 | 7 |
| 3/4 | 5 |
| 3/5 | 4 |
| 3/6 | 1 |
| 4/2 | 2 |
| 4/3 | 5 |
| 4/4 | 2 |
| 4/5 | 5 |
| 4/6 | 1 |
| 4/7 | 1 |

| | |
|---|---|
| 5/4 | 2 |
| 5/5 | 1 |

**Table S5**. FISH on metaphase spreads from matched patient derived xenographs in 2 replicates

| **Number of signals** | **Probe 651L9 (Replicate 1)** | **Probe 651L9 (Replicate 2)** |
|---|---|---|
| 1 | 11 | 4 |
| 2 | 28 | 22 |
| 3 | 11 | 18 |
| 4 | 12 | 15 |
| 5 | 0 | 6 |
| 6 | 2 | 4 |
| 7 | 0 | 1 |
| Clusters* | 36 | 30 |

*Clusters of signals close to each other

**Table S6.** Templated insertion threads identified in PCAWG. (TableS6_pcawg.templated.insertion.threads.xlsx)

**Table S7.** Overview of expression effects, methylation effects, and genetic variants between samples and between haplotypes. (TableS7_overview_gen.meth.expr.xlsx)

**Table S8**. Allele specific expression, allele specific methylation, and allele specific genomic copy number of primary tumor sample. (TableS8_asm_ase_cn_primary_tumor.xlsx)

**Table S9.** Overview of the leafcutter analysis on BASP1. Shown are the splicing cluster overlapping the BASP1 gene and junction usage across the splice cluster. For reference the ALS data (RSP151960) is also shown. (TableS9_splicing_BASP1.xlsx)
